# Altered gut microbial functional pathways in people with Irritable Bowel Syndrome enable precision health insights

**DOI:** 10.1101/2024.02.15.580548

**Authors:** Eric Patridge, Anmol Gorakshakar, Matthew M. Molusky, Oyetunji Ogundijo, Cristina Julian, Lan Hu, Grant Antoine, Momchilo Vuyisich, Robert Wohlman, Guruduth Banavar

## Abstract

Functional gastrointestinal disorders present diagnostic and therapeutic challenges, and there is a strong need for molecular markers that enable early health insights and intervention. Herein, we present an approach to assess the gut microbiome with stool-based gut metatranscriptome data from a large adult human population (*n* = 80,570), using irritable bowel syndrome as an example that features both an abnormal gut microbiome and a spectrum of distinct conditions. We develop a suite of eight gut microbial functional pathway scores, each of which represents the activity of a set of interacting microbial functional features (based on KEGG orthology) relevant to known gut biochemical activities. We use a normative approach within a subpopulation (*n* = 9,350) to define “Good” and “Not Optimal” activities for these transcriptome-based gut pathway scores. We hypothesize that Not Optimal scores are associated with irritable bowel syndrome (IBS) and its subtypes (i.e., IBS-Constipation, IBS-Diarrhea, IBS-Mixed Type). We show that Not Optimal functional pathway scores are associated with higher odds of IBS or its subtypes within an independent cohort (*n* = 71,220) using both the Rome IV Diagnostic Questionnaire as well as self-reported phenotypes. Rather than waiting to diagnose IBS after symptoms appear, these functional pathway scores can help to provide early health insights into molecular pathways that may contribute to IBS. These molecular endpoints could also assist with measuring the efficacy of practical interventions, developing related algorithms, providing personalized nutritional recommendations, diagnostic support, and treatments for gastrointestinal disorders like IBS.

**Graphical Abstract:** 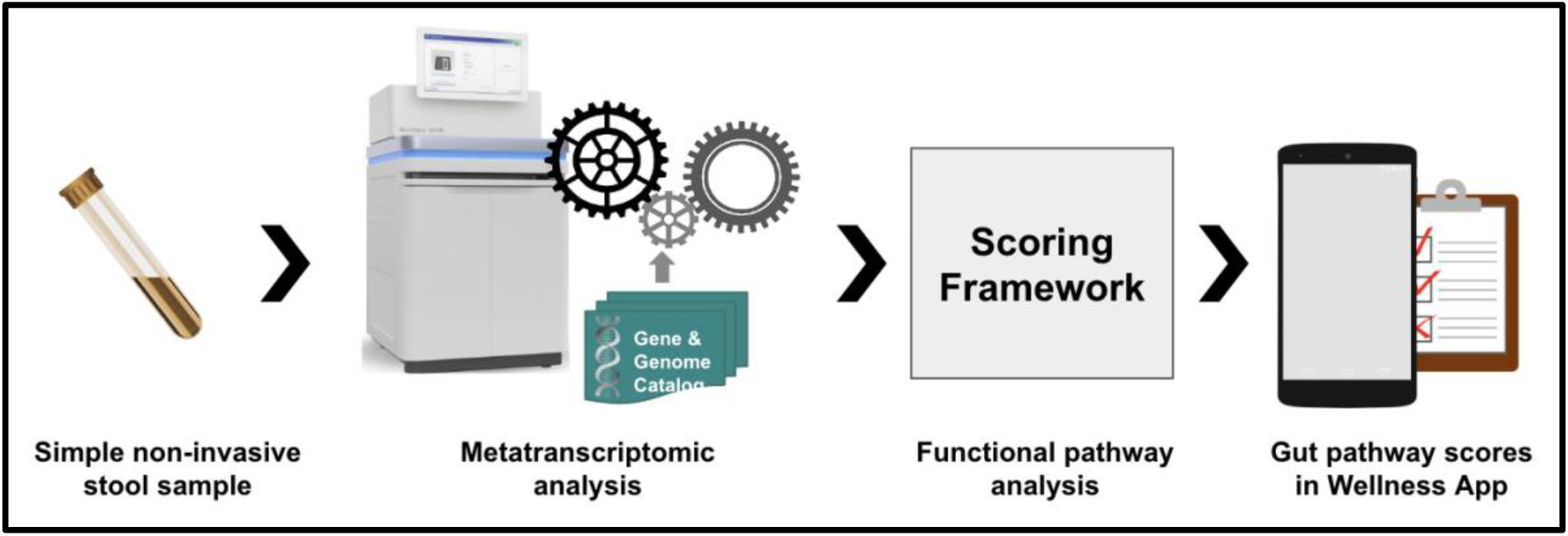

## Introduction

Disorders of gut-brain interactions (DGBIs), previously known as functional gastrointestinal diseases (1), impact nearly 40% of the world’s population (2). According to the 2016 Rome IV criteria, there are more than 33 adult and 20 pediatric DGBIs (3), and there is frequent symptomatic overlap between the DGBI subtypes (4). Their management is complex because known symptoms are subjective and have both varied presentations (5) and varied treatment strategies (6).

Clinically, DGBIs are situated at the interface of the gut-brain connection. They are often chronic, recurrent, debilitating, and do not have an identifiable underlying pathology (7). However, DGBIs are affected by factors which are biological, psychological, and sociological, including sources which are environmental or dietary (8). Common symptoms include altered stool quality, altered motility and gut transit time, hypersensitivity, altered mucosal function, immune disturbances, and altered central nervous system processing (3). One of the most common DGBIs is Irritable Bowel Syndrome (IBS), affecting nearly 11% of the global population (9). Symptoms of IBS vary between patients and may include abdominal pain and discomfort related to abnormal bowel habits such as bloating or distention, along with constipation (for IBS-C) or diarrhea (for IBS-D) – or symptoms of both (for IBS-M) – and the locations and patterns of these may change over time (10). (It should be noticed that IBS-M is a distinct category and not simply an aggregation of IBS-C and IBS-D.) Further nuances of the symptoms may also distinguish “stool consistency”, “gut transit time”, and “motility”; for example, with IBS-C, it is possible to experience an overall slower gut transit time with increased motility localized upstream of a blockage (i.e., the small intestines), and the associated bowel movement could have either increased or decreased firmness, depending on the stool consistency itself (11,12). As a result of the range and variability of overlapping symptoms, patients with DGBIs like IBS often present diagnostic and therapeutic challenges for healthcare providers. For many patients, IBS is a diagnosis of exclusion, and this forces patients to pursue a barrage of medical procedures in order to exclude other diagnoses, while experts plead for easier ways to identify and address the classical symptoms (13,14).

Since IBS is associated with an altered microbiome (15), it may be possible to leverage the gut microbiome for health insights and treatment strategies (16). At the same time, it is important to consider that IBS is also a composite of multiple IBS subtypes (i.e., IBS-Constipation [IBS-C], IBS-Diarrhea [IBS-D], IBS-Mixed Type [IBS-M]), and each subtype may have a distinct microbiome signature (17,18). Typical research efforts focus on profiling metabolites and/or species from the microbiome as part of mechanistic research focused on identifying the underlying causes of IBS (19). Epidemiological reports further advocate for data-driven insight into host, microbiome, and dietary interactions, and multi-omic efforts have already demonstrated some ability for metatranscriptomics to differentiate IBS subtypes (20,21). Still, there is no consensus on a set of biomarkers, microbial functions, or species associated with IBS or its subtypes (15,22). Furthermore, efforts to understand or treat IBS primarily focus on molecular-based diagnostics (23), which are typically reactive measures (24–26), and there remain few options which individuals or providers can use in proactive healthcare (27). Considering the significant challenges and costs associated with identifying and treating IBS and other DGBIs (14,28,29), there is ample opportunity to augment healthcare services with wellness tools.

While the world waits for diagnostics, mechanistic elucidations, and preventative treatments, one possible step to advance healthcare for DGBIs like IBS is by leveraging the altered gut microbiome for early health insights. Historically, chronic diseases such as IBS or other DGBIs are often linked to altered microbiomes with particular focus on taxonomic representations (i.e., species, genus, phylum, etc.), but in many cases, the active functions of those microbes are far more important to human health than are taxonomic representation (30–32). Recently, we developed a stool-based test which measures microbial functions (i.e., gene expression) in stool samples and quantifies them using functions defined by the KEGG Orthology (33,34) during high-resolution metatranscriptome sequencing of the gut microbiome (Figure 1). Insights about the gut microbiome are delivered through a series of gut wellness scores, which then inform downstream personalization through our precision wellness application (35). Ultimately, these scores offer valuable early health insights that can contribute to the development of therapeutic interventions to modulate microbial activities, thereby providing a personalized approach to treating DGIBs and promoting improved health outcomes.

**Figure 1.**
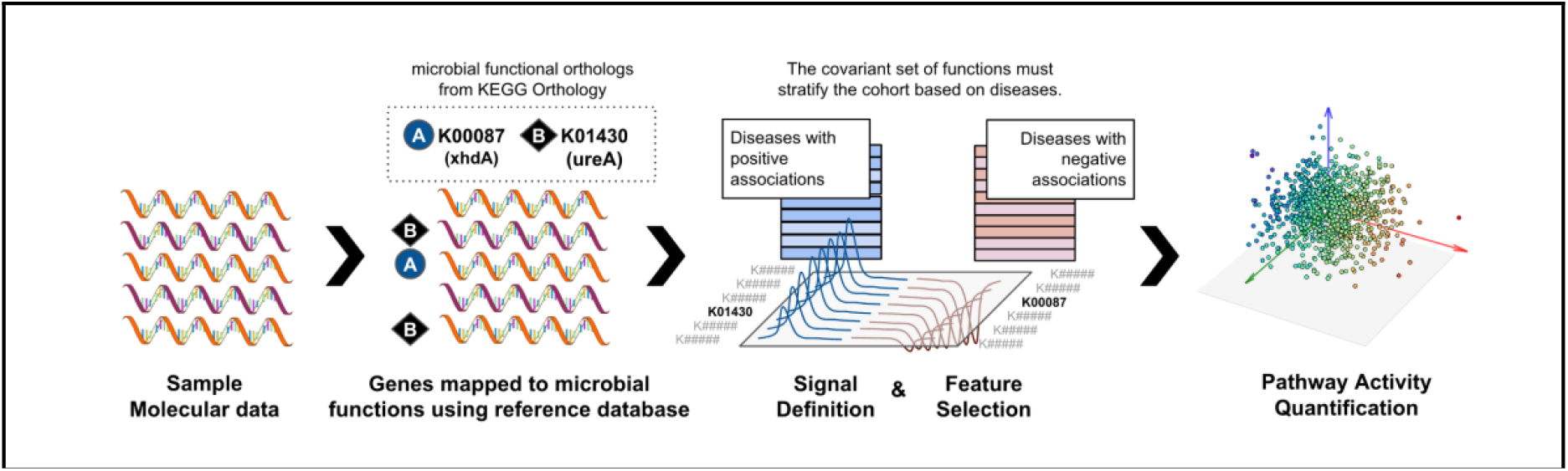
Scoring microbial functions in samples. A suite of scores have been designed for early health insights, and herein, we disclose the *UricAcidProductionPathways* score that includes KEGG identifiers like K00087 and K01430 (Tables 7–8). Signal definition, Feature selection, and Pathway activity quantification are, respectively, Steps 3, 4, and 5 of the Score Development Framework in our earlier publication (34)].

Herein, we demonstrate a scalable approach to quantify the transcriptomic activity of microbial functions from the gut metatranscriptome for applications in health and wellness. We present eight gut pathway scores which were designed to provide insights for both gastrointestinal and systemic health issues. Each score was developed using a large adult population (*n* = 9,350) and a computational method previously demonstrated with saliva (36). The specific purpose of this manuscript is to validate the ability for these transcriptome-based gut pathway scores to assess altered expression levels of microbial functions in stool within an independent cohort (*n* = 71,220). For the current validation, we demonstrate the relationship of the eight gut scores among individuals with IBS and its subtype classifications (IBS-C, IBS-D, and IBS-M), whether cases are defined by self-reported IBS phenotypes or by the Rome IV Diagnostic Questionnaire (37), within an unseen cohort distinct from that used for score development. Each score can be used to identify an altered microbiome, which informs the opportunity to proactively address the altered microbiome.

## Materials and Methods

Stool samples and questionnaire responses were obtained from a general adult population of Viome customers who were at least 18 years old at the time of sample collection and residing in the USA. All customers were informed, and they consented to their data being used for research purposes, as part of the sign-up process for Viome services. At Viome Life Sciences, we assess each research project to determine whether it needs to be reviewed by an Institutional Review Board (IRB). Based on the US Health and Human Services CFR 46.104, section 4(II), the Ethics Program Director determined that this research is exempt from IRB review. This research project solely uses retrospective data from Viome Life Sciences customers. All study data are de-identified; data analysis team members have no access to personally identifiable information. (Clinical trial number: not applicable.)

Study participants collected their stool samples using Viome’s Full Body Intelligence kits, which they received at their homes. Each kit includes a stool collection tube featuring: 1) an integrated scoop to facilitate stool collection, 2) RNA Preservation Buffer (RPB) to maintain the integrity of all RNA molecules during collection and transportation, and 3) sterile glass beads to facilitate sample homogenization. Also included in the kit is a flushable stool sample collection paper, which is placed across the toilet and on which stool is deposited. From the stool, a pea-sized sample is placed inside the collection tube, then vigorously shaken with the RPB to: 1) homogenize the stool, and 2) expose the sample to RPB. As previously published, the RPB has been clinically validated to preserve RNA for up to 28 days at room temperature, and all molecular biology steps were performed in a CLIA-certified laboratory using our clinically validated methodology (35). The method includes DNase treatment, non-informative RNA depletion, cDNA synthesis, size selection, and limited cycles of PCR for adding dual unique barcodes to each sample. All samples were sequenced using the Illumina NovaSeq 6000 platform with 2×150 paired-end read chemistry.

When collecting their stool samples, individuals also respond to an extensive questionnaire allowing them to detail their lifestyle, health history, and health conditions. Lifestyle questions relevant to the current study include use of proton-pump inhibitor (PPI) medication, recent use of antibiotics, and use of disease-specific medication (i.e., antidiarrheals or laxatives/stool softeners). Health history is also collected, resulting in hundreds of self-reported phenotypes which include IBS, IBS-C, IBS-D, and IBS-M, as well as all the disease phenotypes listed in Supplementary Table 1. Symptom and condition questions relevant to the current study include abdominal pain, bloating, stool classification according to the Bristol Stool Form Scale (BSFS), and all questions required for the Rome IV Questionnaire (37–40).

Briefly, the BSFS is an ordinal scale of stool consistency ranging from the hardest (Type 1) to the softest (Type 7). Types 1 and 2 are considered to be abnormally hard stools (indicative of decreased gut transit time (39)) while Types 6 and 7 are considered abnormally loose/liquid stools (indicative of increased gut transit time (39)). Type 3, 4 and 5 are therefore generally considered to be the most ‘normal’ stool form (indicative of average gut transit time (39)).

The Rome IV Diagnostic Questionnaire consists of six questions, beginning with “In the last 3 months, how often did you have pain anywhere in your abdomen?” The answers to these questions are scored and yield a classification of “IBS-C”, “IBS-D”, “IBS-M”, “IBS-U” (for unclassified), or “FALSE” for individuals who do not meet the criteria for any IBS classification (37,41).

### Cohort Development

To deliver health insights from stool samples, we design and validate wellness scores in the context of two cohorts representative of the general adult population. The “development” and “validation” cohorts began with approximately 10,000 and 80,000 samples, respectively. We further exclude outliers based on age, birth sex, and body mass index (BMI). Additional criteria are applied to mitigate artifacts linked to sequencing depth or data sparseness within each cohort. The final score development cohort consists of 9,350 stool samples from adults (59.4% female; Table 1). The final independent validation cohort consists of 71,220 stool samples from adults (66.5% female; Table 1).

**Table 1.** Sociodemographic, anthropometric, medication use, and IBS-related characteristics of the cohorts.

|  |  |  |  |  |
| --- | --- | --- | --- | --- |
|  | Score Development Cohort<br>(*n* = 9,350 stool samples) |  | Independent Validation Cohort<br>(*n* = 71,220 stool samples) |  |
| Female | 59.4% |  | 66.5% |  |
| Age (years) | 46.2 ± 13.0 |  | 44.5 ± 13.6 |  |
| BMI (kg/m²) | 26.1 ± 5.4 |  | 25.8 ± 5.3 |  |
| Ethnicity | 74.2% White<br>From the remaining 25.8%:<br>28.9% Hispanic or Latino<br>28.1% Multi-ethnic<br>19.2% Asian<br>11.1% Black or African American<br>5.6% Arab or Middle Eastern<br>7.0% Other |  | 74.1% White<br>From the remaining 25.9%:<br>28.7% Hispanic or Latino<br>27.4% Multi-ethnic<br>18.5% Asian<br>12.2% Black or African American<br>5.9% Arab or Middle Eastern<br>7.4% Other |  |
| IBS Prevalence | (*self-reported*) | (*Rome IV*) | (*self-reported*) | (*Rome IV*) |
| No IBS | 93.6% | 81.2% | 91.1% | 74.5% |
| Any IBS | 6.4% | 10.0% | 8.9% | 12.1% |
| IBS-C | 1.4% | 2.2% | 2.4% | 2.7% |
| IBS-D | 1.1% | 3.3% | 1.4% | 4.1% |
| IBS-M | 0.9% | 3.9% | 1.5% | 4.6% |
| IBS-U | – | 0.6% | – | 0.6% |
| PPI use<sup>1</sup> | 4.2% |  | 4.3% |  |
| Antibiotic use<sup>2</sup> | 15.4% |  | 13.1% |  |
| Medication use | 0.5% Antidiarrheal meds<br>2.4% Laxatives/stool softeners |  | 0.3% Antidiarrheal meds<br>3.1% Laxatives/stool softeners |  |
| Sample BSFS<sup>3</sup> |  |  |  |  |
| < 3 | 19.4% |  | 19.6% |  |
| 3 < 6 | 52.6% |  | 51.8% |  |
| ≥ 6 | 28.0% |  | 28.7% |  |
| No answer | 0.02% |  | 0.1% |  |
| Abdominal Pain |  |  |  |  |
| Never | 42.4% |  | 37.3% |  |
| < 1 day/wk | 35.8% |  | 36.0% |  |
| 1-6 days/wk | 18.4% |  | 22.7% |  |
| 7 days/wk | 3.3% |  | 3.9% |  |
| No answer | 0.2% |  | 0.3% |  |
| Bloating |  |  |  |  |
| Never | 16.0% |  | 12.7% |  |
| Rarely | 25.8% |  | 22.1% |  |
| Sometimes | 30.6% |  | 30.5% |  |
| Often | 20.0% |  | 24.3% |  |
| Always | 7.5% |  | 10.1% |  |
| No answer | 0.2% |  | 0.3% |  |
| <sup>1</sup> PPI = Self-reported proton-pump inhibitor use. |  |  |  |  |
| <sup>2</sup> Self-reported antibiotics, “currently taking them”, or “have taken them in the past month”. |  |  |  |  |
| <sup>3</sup> Bristol Stool Form Score (average of multiple selection, rounded down). < 3: indicative of decreased gut transit time (1 = Separate hard pellets; 2 = Clumpy log shaped); 3 - 6: indicative of average gut transit time (3 = Log shape with cracks on the surface; 4 = Smooth snake shape; 5 = Soft chunks); ≥ 6: indicative of increased gut transit time (6 = Paste consistency with ragged edges; 7 = Liquid consistency with no solid pieces) |  |  |  |  |

### Metatranscriptomic Analysis

Stool samples are collected and analyzed from individuals who undergo a minimum 8-hour fasting period. With established metatranscriptomic techniques (42), RNA molecules from each stool sample are sequenced. Sequencing reads are aligned to the reference as follows: Viome utilizes a custom reference catalog that encompasses 32,599 microbial genomes from NCBI RefSeq release 205 “complete genome” category, as well as 4,644 representative human gut genomes of UHGG (43), ribosomal RNA (rRNA) sequences, and the human genome GRCh38 (44). Altogether, the custom catalog spans archaea, bacteria, fungi, protozoa, phages, viruses, and the human host, with a total of 98,527,909 microbial genes. To annotate microbial gene functions, the KEGG Orthology (33,34) is adopted, and KEGG orthologs (KOs) are assigned via eggNOG-mapper (45). Taxonomy classification (at all taxonomic levels) and quantification are accomplished using Centrifuge (46). Any reads mapped to the human genome or rRNA sequences are excluded from further analysis but tracked for monitoring. To estimate expression level (or activity) in the sample, an Expectation-Maximization (EM) algorithm (47) is applied to mapped reads. The abundance of respective taxonomic ranks (strains, species, genera, etc.) can be aggregated from the microbial genomes. For the current study, species activity is used in the downstream analyses. The reads mapped to microbial genomes are further extracted and mapped to microbial genes for molecular function or KO quantification. The identified KOs from stool samples are used in downstream analyses, such as score development, validation, and ultimately scoring new samples.

### Score Development

Our stool-based scores are intended to deliver early insights using only microbial functions (i.e., gene expression) from the gut microbiome. We follow a similar approach for score development as we previously reported for saliva-based scores with the oral microbiome (36). In developing these stool-based scores, we explore domain concepts including carbon and energy metabolism, aerobic/anaerobic metabolism, activities related to vitamins and protein cofactors, metabolism of amino acids and nucleic acids, environmental stressors, known pathogenic components, etc. The microbial KOs included in the selection process are identified because of their association with one or more of these physiological processes. During our development process, we leverage the covariance of the selected microbial KOs within a normalized expression dataset, and we aggregate the computationally determined weights, based on the first component (PC1) of Principal Component Analysis (PCA). When the scores are computed on a large cohort, they exhibit a Gaussian-like curve which represents the stratification of health insights and distribution of scores across the population (shown in Supplementary Figure 1).

Our five-step process for score development was previously explained in detail (36), including: 1) Domain exploration; 2) Metadata curation; 3) Signal definition; 4) Feature selection; and 5) Pathway activity quantification. The complete details of each step are available in our earlier report but for context are briefly reviewed. Through domain exploration, we define the concepts applicable to the score as well as the entire set of microbial KOs relevant to both those concepts and the score. The metadata curation process includes identifying and creating case/control labels which enable downstream steps as well as the validation presented in this manuscript. Signal definition is the early draft of a score, meaning a small set of KOs demonstrating covariance which begins to distinguish those with comorbidities from those defined as “healthy” within the development cohort. The feature selection process is extremely iterative and serves to strengthen the score’s ability to stratify health insights in a manner consistent with domain knowledge. The final step, pathway activity quantification, is the aggregate quantification of the entire set of KO features in a score by combining the expression levels of the selected KOs. As we previously reported (36), “we derive a pathway score as a weighted function Score = *C1F1* + *C2F2* + … + *CnFn*, where *Fi* is the expression level of the feature and *Ci* is its weight.”

No clinical standards are yet available for transcriptome-based assessments, so the implementation of our novel transcriptome-based scores follows a normative approach (48,49,50). In this normative approach, the development cohort serves as a reference population which is used to define population-level thresholds for the utility of categorizing deviations from the norm for descriptive purposes. Each score computed with these weightings provides a snapshot of the sample’s RNA expression activity with respect to the development cohort. After using the development cohort to design and compute the scores with a numeric value between 0 and 100, the numerical values are further assigned one of the categories, “Good”, “Average”, or “Not Optimal”, based on the score distribution in the development cohort. In the Results section (*Score Development* sub-section), we fully disclose one of these scores and further elaborate on what constitutes “Good” versus “Not Optimal.” We adopt MAD (median absolute deviation), a measure of dispersion and robust to outliers. In order to capture the norm within the population, categorical thresholds are defined using *median* ± 0.75 * *MAD* of the respective score distribution in the development cohort. “Good” refers to the higher end, “Not Optimal” the lower end, and “Average” the middle range.

### Score Validation

An independent validation cohort, consisting of 71,220 stool samples, is used to confirm the reproducibility of score performance. For the validation process, we take a generalized approach and evaluate the ability for each of the eight scores to differentiate gastrointestinal diseases known to be associated with the gut microbiome.

The focus of the validation process in this paper employs odds ratios (ORs) to test how scores perform, using IBS and each subtype (IBS-C, IBS-D, IBS-M) as a spectrum of distinct but related conditions. Cases for each self-reported IBS phenotype were defined by responses to the question: “Please list all of the illnesses you are currently suffering from or diagnosed with.” The list included IBS, IBS-C, IBS-D, and IBS-M, (with “IBS” as the composite of all IBS subtypes) and all such metadata labels defined in the development cohort are also applied to the validation cohort. Cases for Rome IV-determined IBS classifications were defined using the independent and clinically validated Rome IV Diagnostic Questionnaire. Controls are carefully curated using to identify and exclude non-IBS phenotypes likely to have a “Not Optimal” score; therefore, control definitions are score-specific (see Supplementary Table 1). Cases are matched to controls (1:5) based on age, birth sex, BMI, and the month when each stool sample was sequenced. Age matching allows for ±5 years with respect to case. For BMI, matching is performed within defined categories: BMI < 15 as an outlier, 15 ≤ BMI < 18.5 (“underweight”), 18.5 ≤ BMI < 24.9 (“healthy”), 24.9 ≤ BMI < 29.9 (“overweight”), and BMI ≥ 29.9 (“obese”). For “month sequenced”, matching was performed within a window of ±3 months.

Across the two cohorts, scores were assigned one of three categories: “Good”, “Average”, or “Not Optimal”; for ORs, we compare the odds for “Not Optimal” scores to the odds for “Good” scores. Using score category (i.e., “Not Optimal” or “Good”) as the independent variable and IBS subtype as the dependent variable, a logistic regression model is applied while adjusting for age, BMI, PPI use or antibiotic use, and other disease-specific confounders. OR is defined as the odds of having a “Not Optimal” score with an IBS subtype over having a “Good” score with the same IBS subtype. Statistical comparison of ORs between matched case/control samples is carried out using the two-tailed Mann-Whitney U test, and the Benjamini-Hochberg correction is applied to control the False Discovery Rate.

Venn diagrams included in supplementary materials were created from separate efforts which employed differential expression of species for cases of IBS-C, IBS-D, or IBS-M, excluding cases with multiple IBS subtypes. Controls were defined as those without any self-reported comorbidities. Whereas ORs accounted for individuals taking PPIs or antibiotics, these were excluded from differential expression analyses for the Venn diagrams included in supplementary materials. Otherwise, methodology for matching was the same process as followed for ORs, where cases are matched to controls (1:5) based on age, birth sex, BMI, and month sequenced. For visualization purposes, differentially expressed species (p-value ≤ 0.05 and log(Fold Change) ≥ 0.6) were aggregated to the genus level.

## Results

### Cohort Characteristics

We employ two large cohorts: 1) a score development cohort is used to facilitate a heuristic approach for feature (KO) selection during our score design process; and 2) an independent validation cohort is used to assess the performance of each score. The composition of both cohorts are comparable; the development cohort (9,350 stool samples from adults; 59.4% female) has an average age of 46.2 ± 13.0 years and BMI of 26.1 ± 5.4 kg/m^2^, while the validation cohort (71,220 stool samples from adults; 66.5% female) has an average age of 44.5 ± 13.6 years and BMI of 25.8 ± 5.3 kg/m^2^.

Table 1 shows characteristics for each cohort at the time of sample collection, including sociodemographics (age, birth sex, ethnicity), anthropometrics (BMI), and medication use (PPI, recent antibiotics, anti-diarrhea meds, and laxatives/stool softeners). Additional characteristics related to IBS include both self-reported and Rome IV-defined IBS phenotypes, the Bristol Stool classification of the bowel movement that was sampled for the cohort, abdominal pain frequency over the 3 months prior to sampling, and frequency of bloating.

Since our validation of stool-based scores focuses on individuals with IBS, we present the IBS-related characteristics across the two cohorts in Tables 2–5. In each table, IBS is defined as the complete set of individuals who self-reported IBS or any IBS subtype (IBS-C, IBS-D, IBS-M). The trends reported are similar across both the score development cohort and the independent validation cohort.

**Table 2.**
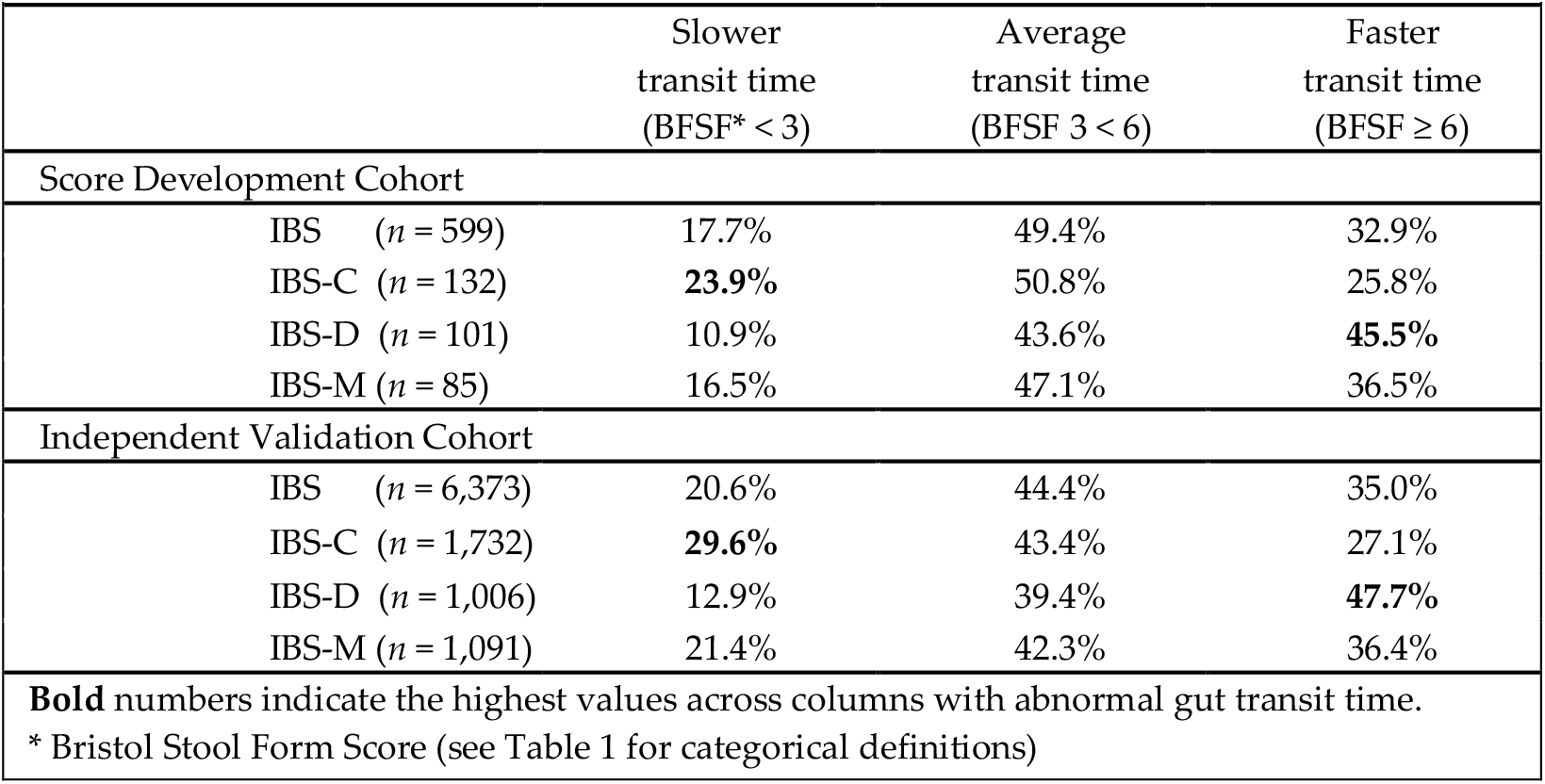
Gut transit time as measured by Bristol Stool Scores for the sampled (analyzed) bowel movement from IBS (any subtype) and IBS subtypes across cohorts (totaling 6,972 individuals). Gut transit times are as expected across subtypes, and the two cohorts have similar characteristics.

Table 2 shows the Bristol Stool classification of the sampled bowel movement, stratified by IBS subtype. Expected trends are highlighted in bold: a higher proportion of individuals with IBS-C self-report slower gut transit times while a higher proportion of those with IBS-D self-report faster gut transit times. Individuals with IBS-M report Bristol Stool classifications similar to the entire set of individuals with IBS, which also closely follows the general cohort population.

Table 3 shows responses of recent abdominal pain, broken down for those with IBS and those without IBS. Expected trends are highlighted in bold: across both cohorts, individuals with IBS reported more frequent abdominal pain than those without IBS. When stratified by IBS subtype, the numbers were generally similar across all subtypes (not shown).

**Table 3.**
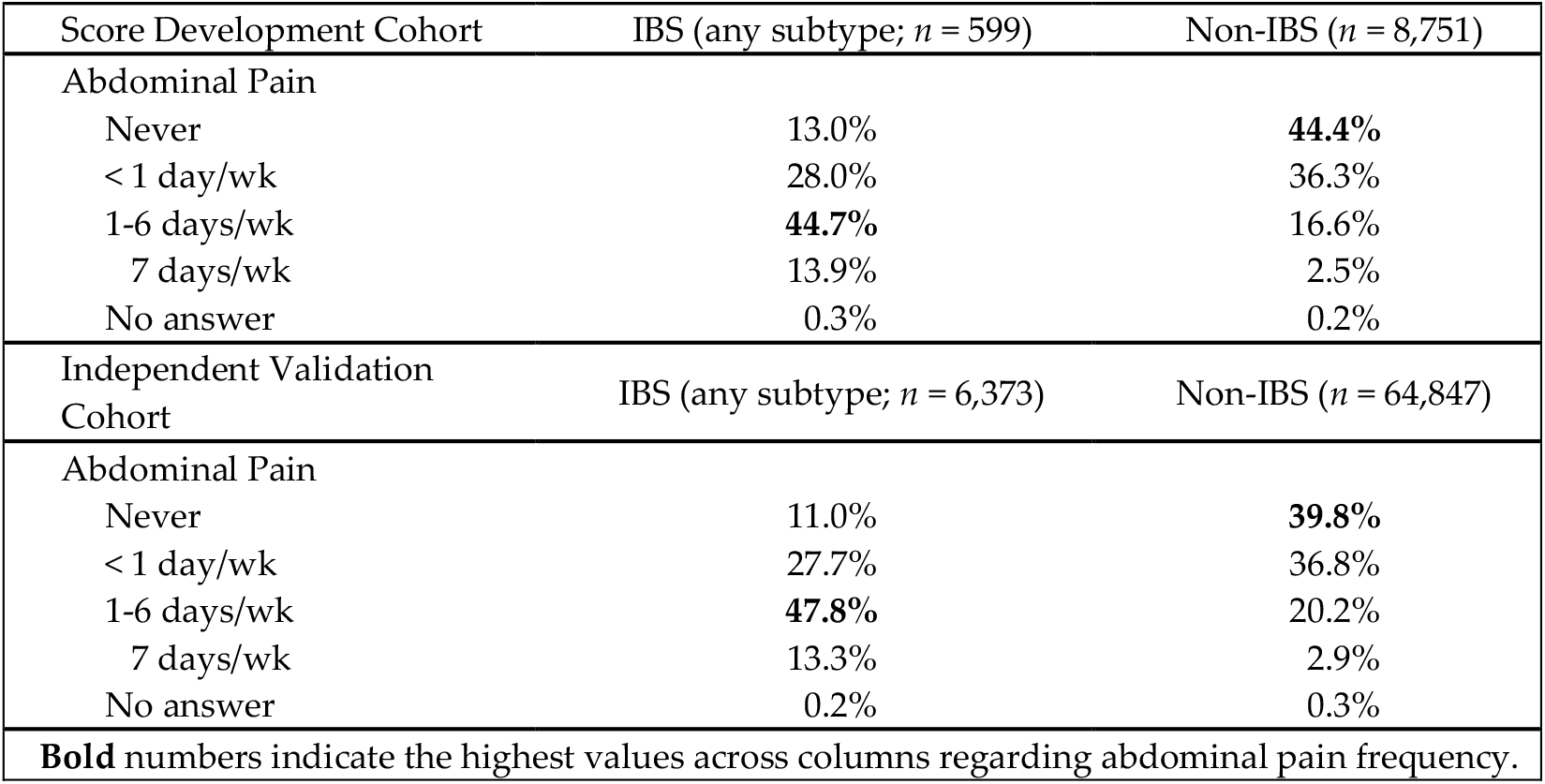
Abdominal pain reported by individuals with IBS (*n* = 6,972) and Non-IBS (*n* = 73,598). Breakdown of abdominal pain is as expected for IBS/Non-IBS, and the two cohorts are similar.

Those individuals who reported any abdominal pain (Table 3) were also asked for the Bristol Stool classification of their recent atypical bowel movements, and these are stratified across IBS subtypes in Table 4. As the numbers show, the trends highlighted in bold in Table 2 are also present in Table 4. A higher proportion of individuals with IBS-C indicated decreased gut transit time (“usually constipation; BSFS 1-2”), while a higher proportion of individuals with IBS-D indicated increased gut transit time (“usually diarrhea; BSFS 6-7”). A higher proportion of individuals with IBS-M indicated mixed gut transit time (“both constipation and diarrhea”).

**Table 4.** Gut transit time as measured by Bristol Stool Scores for recent atypical bowel movements from IBS (any subtype) and IBS subtypes across cohorts (totaling 6,972 individuals). Breakdown of gut transit time pain is as expected across subtypes, and the two cohorts are similar.

|  |  | Slower<br>transit time<br>(BFSF** < 3) | Average<br>transit time<br>(BFSF 3 < 6) | Faster<br>transit time<br>(BFSF ≥ 6) | Mixed<br>transit time<br>(BFSF < 3 & ≥ 6) | No<br>answer* |
| --- | --- | --- | --- | --- | --- | --- |
| **Score Development Cohort** |  |  |  |  |  |  |
| IBS | (*n* = 599) | 21.7% | 4.8% | 28.7% | 31.4% | 13.4% |
| IBS-C | (*n* = 132) | **43.9%** | 3.8% | 6.1% | 28.8% | 17.4% |
| IBS-D | (*n* = 101) | 2.0% | 0.0% | **67.3%** | 18.8% | 11.9% |
| IBS-M | (*n* = 85) | 11.8% | 2.4% | 29.4% | **49.4%** | 7.1% |
| **Independent Validation Cohort** |  |  |  |  |  |  |
| IBS | (*n* = 6,373) | 22.3% | 5.0% | 30.0% | 31.5% | 11.2% |
| IBS-C | (*n* = 1,732) | **50.3%** | 4.1% | 6.6% | 26.0% | 13.0% |
| IBS-D | (*n* = 1,006) | 2.7% | 1.8% | **65.2%** | 15.8% | 14.5% |
| IBS-M | (*n* = 1,091) | 9.4% | 3.3% | 24.4% | **53.7%** | 9.2% |

**Bold** numbers indicate the highest values across columns with abnormal gut transit time.
\* Responses were not recorded for individuals who never experienced abdominal pain.
\*\* Bristol Stool Form Score (see Table 1 for categorical definitions)

Table 5 shows responses of typical bloating, broken down for those with IBS and those without IBS; across both cohorts, individuals with IBS reported more frequent bloating than those without IBS. When stratified by IBS subtype, the numbers were generally similar across all subtypes (not shown).

**Table 5.**
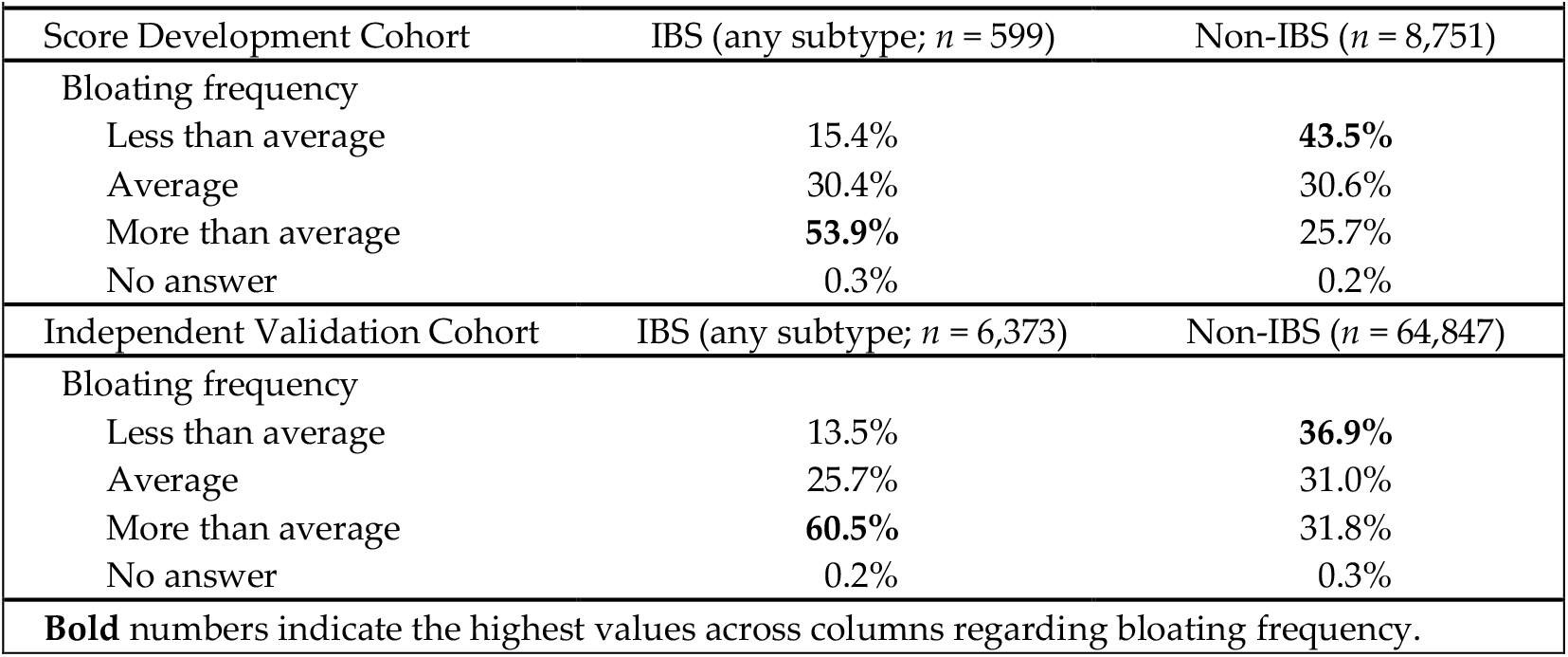
Bloating frequency reported by individuals with IBS (*n* = 6,972) and Non-IBS (*n* = 73,598). Bloating is as expected for IBS/Non-IBS, and the two cohorts have similar characteristics.

|  |  |  |
| --- | --- | --- |
| Score Development Cohort | IBS (any subtype; $n = 599$) | Non-IBS ($n = 8,751$) |
| Bloating frequency |  |  |
| Less than average | 15.4% | **43.5%** |
| Average | 30.4% | 30.6% |
| More than average | **53.9%** | 25.7% |
| No answer | 0.3% | 0.2% |
| Independent Validation Cohort | IBS (any subtype; $n = 6,373$) | Non-IBS ($n = 64,847$) |
| Bloating frequency |  |  |
| Less than average | 13.5% | **36.9%** |
| Average | 25.7% | 31.0% |
| More than average | **60.5%** | 31.8% |
| No answer | 0.2% | 0.3% |
| **Bold numbers indicate the highest values across columns regarding bloating frequency.** |  |  |

### Metatranscriptomic Analysis

We present high-level metrics of development cohort (*n* = 9,350) and independent validation cohort (*n* = 71,220) to demonstrate sample data quality. On average, each sample has 10.6 ± 4.28 million reads from paired-end fastq files when combined across both cohorts. The average KO richness or distinct KOs associated with each sample for development and validation cohorts combined is 3,641 ± 699. Species richness, defined as distinct species identified in each sample, across development and validation cohorts combined, is 1,451 ± 518. Effective sequencing depth, defined as the total reads mapped to a recognized element like KO or species for both cohorts combined, is 3.32 ± 2.20 million.

### Score Development

Our Score Development Framework is a 5-step process (36), and herein, we use *UricAcidProductionPathways* as an example while we talk through each step. A high-level summary for each of the eight gut pathway scores appears in Table 6, including key functions and a brief description of microbial activities. Altogether, the eight scores comprise a total of 278 KOs (259 distinct KOs), where 242 KOs (or 93%) appear only once across all scores and 17 KOs (or 6.5%) each appear in 2-3 scores. Any two scores share a maximum of three KOs. The number of KOs in each score ranges from 19 to 63, with the final number determined heuristically rather than stringent limits. While the process is similar for each score, all scores are developed independently from each other, with key functions, components, metadata and phenotype labels driving feature selection.

**Table 6.** High level overview of Stool Pathways Scores.

| **Table 6.** High level overview of Stool Pathways Scores |  |  |  |
| --- | --- | --- | --- |
| **Score Name** | **Key Functions** | **Microbial Activities** | **KOs** |
| *ButyrateProductionPathways* | Butanoate metabolism<br>Lysine degradation<br>Fatty acid degradation<br>BCAA<sup>1</sup> degradation<br>Propanoate metabolism | Accumulation of beneficial butyrate from pyruvate and by degradation of fatty acids, lysine, and BCAAs. | 39 |
| *GABAProductionPathways* | Butanoate metabolism<br>Pentose/glucuronate interconversions<br>Vitamin B6 metabolism<br>Amino/nucleotide sugar metabolism | Accumulation of GABA<sup>2</sup> from various substrates including butanoate. Along with synthesis of cofactors needed for enzymes that produce GABA, including vitamin B6. | 63 |
| *LacticAcidProductionPathways* | Lactate transport<br>Pyruvate metabolism<br>Other glycan degradation | Metabolism and transport of butyrate, pyruvate, and lactate, including carbon flow specific to the accumulation of lactic acid. | 30 |
| *LPSBiosynthesisPathways* | Lipopolysaccharide biosynthesis<br>KDO2-lipid A biosynthesis<br>Nucleotide sugar biosynthesis | Accumulation of beneficial and deleterious LPS modifications, nucleotide sugars, and related processing of glutamine/glutamate/ammonia. | 24 |
| *MethanogenesisPathways* | Methane metabolism<br>Formaldehyde assimilation<br>Fructose/mannose metabolism<br>Pentose phosphate pathway | Several methods of methane production, ammonia release from amino acids, and pentose phosphate activity. | 47 |
| *OxalateMetabolismPathways* | Oxalate degradation<br>Glyoxylate/dicarboxylate metabolism<br>Ascorbate/aldarate metabolism | Accumulation of oxalate and glyoxylate precursors, including the metabolism of related vitamins and carbon compounds. | 28 |
| *SulfideProductionPathways* | Sulfur metabolism<br>Taurine/hypotaurine metabolism<br>Glycine/serine/threonine metabolism | Tricarboxylic acid-related energy processes as well as the generation of sulfide from organic sulfur-containing amino acids and inorganic sources like sulfite and sulfate. | 28 |
| *UricAcidProductionPathways* | Nucleotide metabolism<br>Purine metabolism<br>Hypoxanthine oxidation<br>C4-Dicarboxylate sensing | Beneficial production of trehalose and processing of purines like adenine, guanine, inosine, xanthine, and urate. | 19 |
| <sup>1</sup> BCAA = Branched-chain amino acid. |  |  |  |
| <sup>2</sup> GABA = Gamma-aminobutyric acid. |  |  |  |

The first step in the framework is domain-driven curation for potential inclusion of features (KOs) in score development. As an example, the *UricAcidProductionPathways* score was designed by curating “key components” and “key functions” centered around microbial nucleotide, nucleoside, and purine degradation generating uric acid (Table 7). KOs were considered for inclusion if they impacted these “in scope” parameters.

**Table 7.** Scope of the *UricAcidProductionPathways* score, which serves as one example from the suite of available wellness scores.

|  |  |  |
| --- | --- | --- |
| **Key components** | Nucleotides | AMP, GMP, 3'-GMP, IMP |
|  | Nucleosides | Adenosine, guanosine, inosine, xanthosine |
|  | Purines | Adenine, guanine, hypoxanthine, xanthine, uric acid |
|  | Enzymes | Oxidoreductases, deaminases, nucleotidases |
|  | Cofactors | NADH, molybdenum, FAD |
| **Key functions** | Purine metabolism | Hypoxanthine oxidation |
|  | Nucleotide metabolism | Adenosine degradation, guanosine degradation |
|  | Energy extraction | Anaerobic fermentation |
|  | Two-component systems | C4-Dicarboxylate sensing and transport |
|  | Other | Molybdenum biosynthesis, quorum sensing |

The second and third steps involve metadata curation and signal definition in our development cohort. These steps provide information useful for step four, feature selection, which involves the iterative refinement of the KOs that define a score. During design of *UricAcidProductionPathways* across steps 2-4, we selected KOs based on their impact on differentiating several comorbidities and other health insights associated with uric acid (i.e., gout, kidney stones, type 2 diabetes) such that the samples labeled with those comorbidities were maximally differentiated from samples without those comorbidities, according to the metric of “mean score difference.”

The final step involves pathway activity quantification. The score definition for *UricAcidProductionPathways* is depicted in Table 8, alongside the corresponding weights (loadings) derived from the first principal component (PC1) of a principal component analysis (PCA) (see Materials and Methods). The normalized expression data for the development cohort (*n* = 9,350) appears in Figure 2A (including colorized PC1), and the loadings from Table 8 are visualized in Figure 2B. The KOs with negative loadings represent microbial uric acid production, which is undesirable and therefore higher expression of these features would result in a lower score. Thus, an overall greater expression of KOs with negative loadings directs a numerical score towards “Not Optimal,” which represents all values below “*median1 −0.75 * MAD*,” with respect to the development cohort. The loadings in Table 8 also include KOs with positive loadings, which represent uric acid degradation or alternative metabolism, and a higher expression of these features would result in a higher score. Thus, an overall greater expression of KOs with positive loadings directs a numerical score towards “Good,” which represents all values above “*median* + 0.75 * *MAD*,” with respect to the development cohort.

**Table 8.** Score design for *UricAcidProductionPathways*. Loadings originate from the first principal component of PCA analysis using the development cohort.

| Loadings | KEGG Ortholog (KO) and Description |
| --- | --- |
| 0.42707 | K07651 (resE); two-component system, OmpR family, sensor histidine kinase |
| 0.30347 | K04076 (lonB); ATP-dependent Lon protease |
| 0.27966 | K18011 (kamE); beta-lysine 5,6-aminomutase beta subunit |
| 0.27711 | K18532 (AK6, FAP7); adenylate kinase |
| 0.24838 | K01236 (treZ, glgZ); maltooligosyltrehalose trehalohydrolase |
| 0.06793 | K01430 (ureA); urease subunit gamma |
| 0.05184 | K00948 (PRPS, prsA); ribose-phosphate pyrophosphokinase |
| 0.04766 | K00839 (pucG); (S)-ureidoglycine-glyoxylate transaminase |
| -0.02885 | K21053 (ade); adenine deaminase |
| -0.06887 | K20967 (MOCS1); GTP 3',8-cyclase / cyclic pyranopterin monophosphate synthase |
| -0.08136 | K00087 (ygeS, xdhA); xanthine dehydrogenase molybdenum-binding subunit |
| -0.10569 | K13479 (ygeT, xdhB); xanthine dehydrogenase FAD-binding subunit |
| -0.18014 | K15973 (mhqR); MarR family transcriptional regulator |
| -0.22229 | K13483 (yagT); xanthine dehydrogenase YagT iron-sulfur-binding subunit |
| -0.22525 | K00364 (guaC), GMPR; GMP reductase |
| -0.26568 | K11711 (dctS); two-component system, LuxR family, sensor histidine kinase |
| -0.26922 | K03787 (surE); 5'/3'-nucleotidase |
| -0.30322 | K07084 (yuiF); putative amino acid transporter |
| -0.32907 | K10558 (lsrA, ego); AI-2 transport system ATP-binding protein |

**Figure 2.**
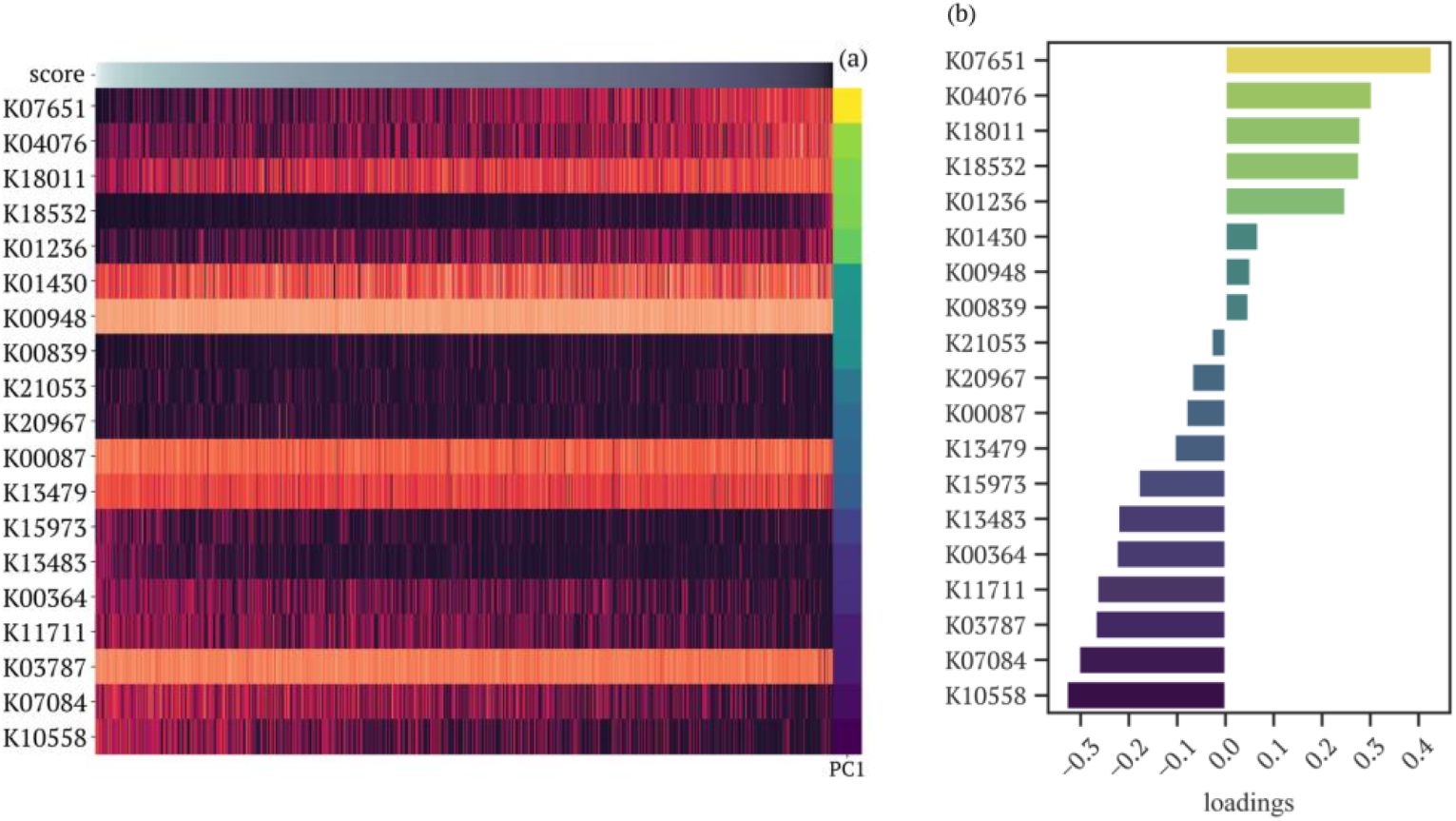
Raw data and loadings from Principal Component Analysis for *UricAcidProduction-Pathways*. **(a)** Normalized expression data from the development cohort (*n* = 9,350) is subsetted by the 19 features for *UricAcidProductionPathways*. All samples are sorted along the x-axis using score values (colorized at top) and all features are sorted along the y-axis using the loadings from Principal Component 1 (PC1; colorized loadings at right). **(b)** Loadings for each of the 19 features of PC1 are visualized for *UricAcidProductionPathways*. Colorization of PC1 in **(a)** is aligned with the loadings in **(b)**.

### Score Validation

We use the independent validation cohort to examine how the eight gut pathway scores behave with IBS and its subtypes; our hypothesis, based on the domain knowledge used to design the score, was that non-optimal scores would associate with IBS and its subtypes. We choose IBS and IBS subtypes (IBS-C, IBS-D, IBS-M) to stratify the scores, because these IBS subtypes have altered gut microbiomes and, to some extent, share underlying symptoms such as altered gut transit, altered stool consistency, abdominal pain, and bloating (51,52).

For simplicity, we only report on “Not Optimal” scores, using “Good” scores as a reference point. The translational interpretation for each “Not Optimal” score is presented in Table 9. For example, in the case of *UricAcidProductionPathways*, a “Not Optimal” score indicates the microbial gene expression contains more functions with negative loadings and fewer functions with positive loadings from Table 8; this may be interpreted as increased uric acid production (Table 9).

Focusing on “Not Optimal” scores, we present ORs for IBS and its subtypes as defined using self-reported IBS phenotypes (Figure 3) or as defined by the independent and clinically validated Rome IV Diagnostic Questionnaire (Figure 4), where the vertical line at 1 represents the reference category (“Good” scores). Numerical values for these are shown in Supplementary Tables 2 and 3, respectively. The ORs indicate the likelihood that someone with a “Not Optimal” score also has the IBS subtype (without informing about causality). As seen in Figures 3 and 4, ORs demonstrate that the gut pathway scores can variably differentiate IBS subtypes. The scores also behave similarly whether IBS and its subtypes are defined using self-reported IBS phenotypes or using the Rome IV Diagnostic Questionnaire.

**Figure 3.**
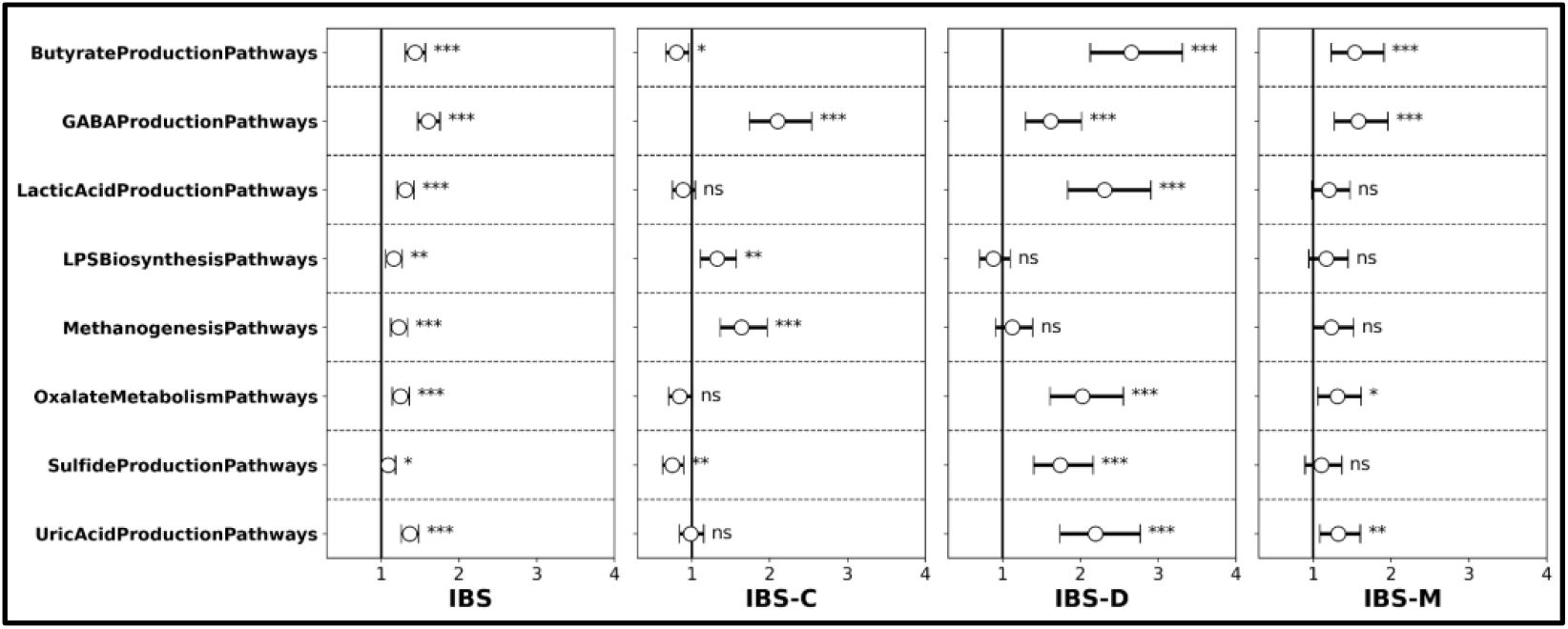
Odds ratios for each “Not Optimal” score stratified with cases defined by self-reported IBS phenotypes. “Good” scores serve as the reference group and are indicated by the vertical line at 1. The maximum number of cases available for 1:5 matching include: 6,373 for IBS; 1,732 for IBS-C; 1,006 for IBS-D; and 1,091 for IBS-M. Definitions for “controls” are score-specific and specified in Extended Data Table 1, including the minimum size of the control pool available for 1:5 matching. p ≤ 0.05 (*); p ≤ 0.01 (**); p ≤ 0.001 (***); not significant (ns). Numerical values for all odds ratios in Extended Data Table 2.

**Figure 4.**
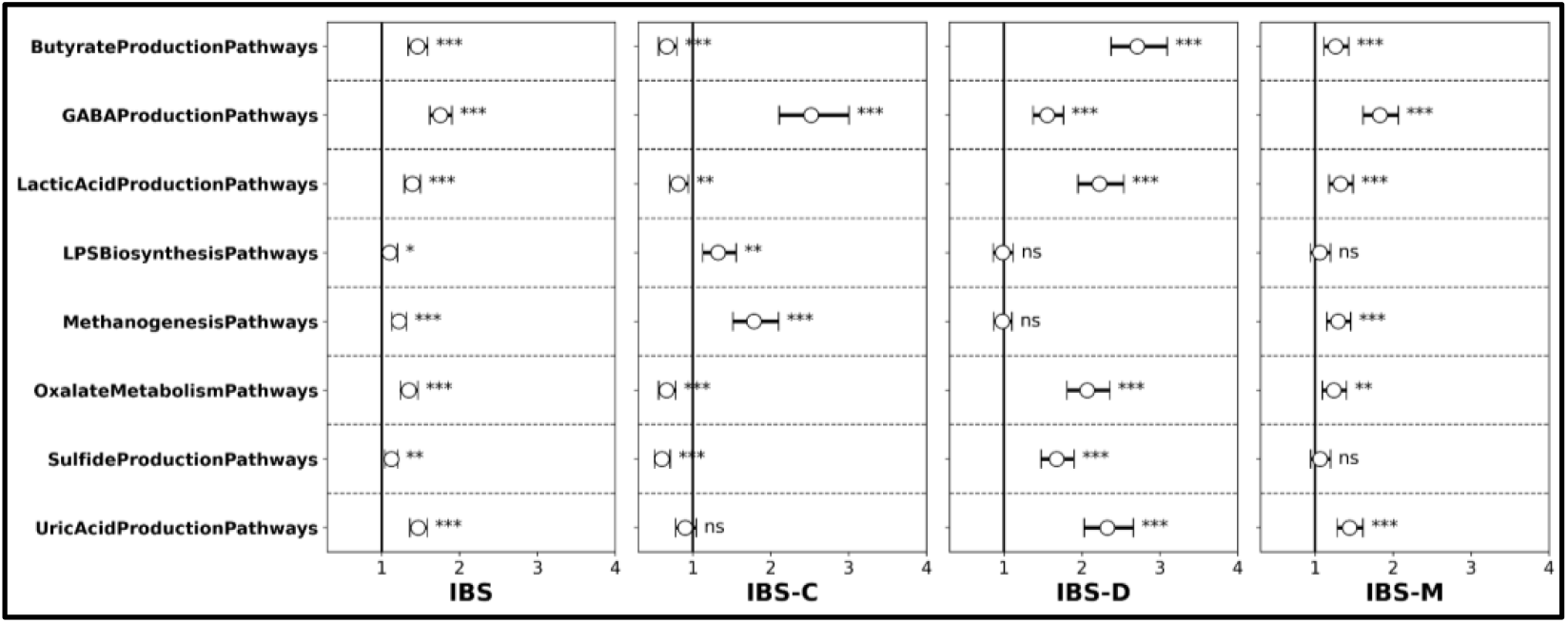
Odds ratios for each “Not Optimal” score stratified with cases defined by Rome IV Diagnostic Questionnaire. “Good” scores serve as the reference group and are indicated by the vertical line at 1. The maximum number of cases available for 1:5 matching include: 8,629 for IBS; 1,942 for IBS-C; 2,952 for IBS-D; and 3,309 for IBS-M. Definitions for “controls” are score-specific and specified in Extended Data Table 1, including the minimum size of the control pool available for 1:5 matching. p ≤ 0.05 (*); p ≤ 0.01 (**); p ≤ 0.001 (***); not significant (ns). Numerical values for all odds ratios are shown in Extended Data Table 3.

Although this manuscript focuses on gut pathway scores, we also make our findings more accessible to the readership by including context for the microbiota (Supplementary Figures 2, 3, 4, and 5). For IBS cases, all significantly increased species are depicted in Supplementary Figure 2 (self-reported IBS phenotypes) or Supplementary Figure 4 (Rome IV-determined IBS phenotypes) while all significantly decreased species are depicted in Supplementary Figure 3 (self-reported IBS phenotypes) or Supplementary Figure 5 (Rome IV-determined IBS phenotypes). All information in the Venn diagrams has been collapsed to the genus level.

## Discussion

Microbial functions within the gut are important for understanding, preventing, and treating many health issues, including DGBIs like IBS. The wellness scores we validate in this manuscript provide early health insights based on microbial functions (KOs) detected through metatranscriptomic analysis of the gut microbiome without focusing on the composition of the microbiota. Rather than serving as diagnostics, the eight gut scores (Table 6) provide generalized insights across both systemic and chronic health issues with particular focus on gastrointestinal conditions. For validation of the scores, we select IBS phenotypes in order to leverage both the underlying symptoms shared across IBS subtypes as well as their divergent characteristics like altered gut transit (41,42). This independent validation demonstrates that gut pathway scores can consistently stratify molecular insights based on signals identified during score development.

### Considerations for score development and cohort definitions

The nature of the independent cohort is important to the validation of gut wellness scores. In order to show consistent functionality of the gut pathway scores, it is important to perform validations with an independent cohort that resembles the development cohort. At the highest level, Table 1 establishes general similarity across the two cohorts while Tables 2–5 show they are comparable in terms of anthropometrics, sociodemographics, medication use (PPIs, antibiotics, anti-diarrheals, and laxatives/stool softeners), and IBS phenotypes (i.e., Rome IV responses, stool quality, abdominal pain, bloating, and gut transit time).

Beyond their similarity, the size of each cohort is also important to the process, because large cohorts facilitate both an array of metadata labels and a large sample count for each definition of “case.” For the current study, most of the cohort metadata labels used to define “case” (IBS phenotypes) and “controls” (Supplementary Table 1) are identified through self-reported questionnaires, which could raise concerns since self-reported health information is sometimes questionable (73,74). However, the ORs based on self-reported IBS phenotypes (Figure 3) are consistent with Figure 4, where cases are defined by the independent and clinically validated Rome IV Diagnostic Questionnaire. Furthermore, Tables 2–5 corroborate the labels for IBS and its subtypes. In Tables 2 and 4, variability in self-reported gut transit times align with expectations for self-reported IBS subtypes; those with slower transit times most often self-report IBS-C, while those with faster transit times most often self-reported IBS-D, and those with mixed atypical bowel movements most often self-reported IBS-M. Similar trends are shown in Tables 3 and 5 for the frequency of abdominal pain and bloating (respectively) between those who self-reported IBS and those who did not (“Non-IBS”). As expected, there was no noticeable trend for abdominal pain and bloating when tabulated across IBS subtypes (not shown). Altogether, the corroborating evidence supports our definitions of “case” and “control” as well as our use of the two cohorts in development and validation of the gut pathway scores.

### Unpacking the UricAcidProductionPathways gut score

To maximize transparency and understandability of the gut pathway scores, we disclose the complete score design for *UricAcidProductionPathways*, and we use the Results section (*Score Development* sub-section) to further elaborate on what constitutes “Good” versus “Not Optimal.” The overall design process remains similar for each score, and we fully detail this process in our earlier report for saliva pathway scores (36).

During score development, we deeply explore the domain knowledge to identify biological phenomena central to each score before finalizing an appropriate set of features to define the score. In the case of *UricAcidProductionPathways*, score KOs are broadly related to the biochemical production of uric acid by the gut microbiome; domain-wise, they are broadly associated with the production or degradation of purines (Tables 7–8), and a “Not Optimal” *UricAcidProductionPathways* score indicates increased levels of uric acid (Table 9). The set of KOs in the final score definition (Table 8) are used to subset the normalized expression data of the entire cohort, and the loadings for each KO represent the feature’s contribution to total RNA variability across the subsetted expression data. Since the selection of each feature is tied to its ability to stratify prioritized phenotypes, like IBS, the subsetted expression data is inherently designed to stratify those phenotypes. In the case of *UricAcidProductionPathways*, feature selection was guided by iteratively maximizing “mean score difference” to distinguish phenotypes related to uric acid (i.e., gout, kidney stones, type 2 diabetes).

A brief interpretation of the ORs in Figures 3 and 4 offers additional context for these gut pathway scores. First, the odds of having a “Not Optimal” *UricAcidProductionPathways* score is significantly higher for those who reported IBS, IBS-D, or IBS-M, but not for those with IBS-C. The clinical implication is that those with IBS, IBS-D, or IBS-M are more likely to have activity related to increased uric acid levels, as measured with the *UricAcidProductionPathways* gut score. Understanding the scientific implications of our findings requires further research, because the relationship between uric acid levels and IBS has not been fully elucidated. However, several studies support a link between purine levels and IBS subtypes (72,75– 77), and there are well-documented associations between high uric acid levels (hyperuricemia) and inflammatory diseases (i.e., gout (78,79), hypertension (80), metabolic syndrome (81), and chronic kidney disease (82)) which are, in turn, associated with IBS or its subtypes (71,83–85). Furthermore, purinergic receptors are implicated in secretomotor reflexes of the gastrointestinal tract (86–88) and are actively pursued as IBS drug targets (89,90).

Finally, it may be helpful to contextualize the attention we give to *UricAcidProductionPathways* and its relationship to IBS. Briefly, we disclose the score with the smallest set of KOs in order to maximize conceptual development and discussion within the paper while minimizing confusion. To be clear, the current study is not intended to produce any diagnostic nor to reveal a novel role for uric acid in relation to IBS. Furthermore, we could have reframed *UricAcidProductionPathways* in context of many other diseases, such as gout, but IBS phenotypes are more common, they traverse a variety of symptoms, and a link between IBS and purines like uric acid has been established.

### Interpreting the gut pathway scores with IBS subtypes

The primary goal for the current study is to validate the ability for transcriptome-based gut pathway scores (based on KEGG functions) to assess altered microbiomes in an independent cohort. To this end, we report thirty-two ORs for the eight gut pathway scores, where individuals with self-reported IBS phenotypes have generally higher likelihood to present “Not Optimal” scores (Figure 3). For twenty-two of the ORs, IBS phenotypes are significantly more likely to present “Not Optimal” scores. Eight other ORs are indistinguishable from “Good”, and for the remaining two ORs, IBS-C is less likely to present “Not Optimal” scores. In addition to Figure 3, we also include ORs with cases defined by the Rome IV Diagnostic Questionnaire (Figure 4). Between the two definitions for IBS phenotypes, the scores behave quite similarly, with the primary differences being the level of significance for four of the values in distinguishing “Good” from “Not Optimal” scores.

A single high-level interpretation of the ORs (Figures 3 and 4) is challenging; conceptually, each score is designed to cover distinct biological phenomena. Furthermore, each OR is computed with its own case/control definitions (Supplementary Tables 1-3) but highlighting specifics may prove useful for identifying future areas of research.

Overall, Figures 3 and 4 indicate those with IBS-D present with more “Not Optimal” scores than those with IBS-C, IBS-M, and even the IBS composite. One explanation for this is that faster gut transit times are associated with an altered microbiome (91) and altered fecal metabolites (92), and it is possible that the gut is impacted to a greater degree with IBS-D. For three of the scores which fit this pattern: “Not Optimal” corresponds to increased production of lactic acid with *LacticAcidProductionPathways*, decreased degradation of oxalates with *OxalateMetabolismPathways*, and increased production of sulfide with *SulfideProductionPathways* (Table 9). These ORs agree with the literature, which indicates that IBS-D and/or diarrhea is associated with increased levels of lactic acid (93,94), oxalates (67,95), and sulfides (66,70,96). Importantly, the immunomodulatory and neuroactive properties of these gut-derived metabolites contribute to key features of IBS-D, including visceral pain, stress sensitivity, immune dysregulation, and epithelial dysfunction. Lactic acid is able to modulate immune responses, heighten visceral pain, and alter stress perception— core features of the gut-brain axis disturbances in IBS-D (97,98). Excessive hydrogen sulfide can be cytotoxic and disrupt mitochondria, impair epithelial barrier integrity, and activate inflammatory pathways (99,100). Oxalate can accumulate and induce oxidative stress, compromise epithelial tight junctions, and trigger inflammatory cascades (101,102). Thus, these metabolites may contribute to the dysbiosis and inflammation aspects of IBS subtypes.

One striking trend is the increased likelihood for each of IBS, IBS-C, IBS-D, and IBS-M to present “Not Optimal” *GABAProductionPathways* scores (Figures 3 and 4), corresponding to decreased production of gamma-aminobutyric acid (GABA) as seen in Table 9. GABA is a major inhibitory neurotransmitter and can be produced by gut microbes (103). One role GABA plays in the gastrointestinal tract is to dampen pain, and consistent with our cohort metadata (Table 3), increased abdominal pain is a cardinal feature of IBS (104). It is possible that reduced GABA impairs gastrointestinal pain processing for IBS and its subtypes (105). Beyond the gut, reduced GABA function is associated with depression and anxiety (106), and IBS patients are at higher risk of developing depression or anxiety compared to the general population (107). Many studies support that GABA dysregulation plays a role in IBS, and this is consistent with our findings.

Of all the ORs, the largest values are for *ButyrateProductionPathways* with IBS-D, for which a “Not Optimal” score corresponds to decreased production of butyrate (Table 9). Thus, individuals with IBS-D may have reduced activity related to butyrate production, as measured with the *ButyrateProductionPathways* gut score. While there are conflicting reports regarding fecal butyrate levels in IBS-D, this condition is generally associated with a reduction in butyrate-producing microbes such as *Faecalibacterium prausnitzii* and *Roseburia* spp (52,54). Of course, the gastrointestinal role of butyrate and other short-chain fatty acids (SCFAs) is quite complex and their specific impact on IBS-D remains to be elucidated. However, butyrate is known to profoundly affect both local intestinal physiology and systemic immune function. In the gut, it serves as a crucial colonic energy source, enhances epithelial barrier integrity, reduces mucosal inflammation, and modulates visceral sensitivity—key factors in IBS-D pathophysiology (108,109). SCFAs like butyrate also influence the gut-brain axis by interacting with enteroendocrine cells, modulating serotonin biosynthesis, and signaling, thereby potentially impacting mood and stress responses commonly comorbid with IBS (110,111). Furthermore, butyrate supplementation has shown an ability to mitigate abdominal pain and improve stool consistency in IBS-D patients (112,113).

Our findings also indicate that those with IBS-C are more likely to present “Not Optimal” *LPSBiosynthesisPathways* scores (Figures 3 and 4), corresponding to increased deleterious microbial LPS production (Table 9). LPS is the major surface membrane component of almost all Gram-negative bacteria (114), and its role in promoting inflammation is well-documented (115). However, there is tremendous heterogeneity in LPS structural modifications (116), and associations between microbes and IBS symptoms have been inconsistent or even contradictory (117). In contrast to expectations, some components and versions of LPS are even antagonistic to inflammatory responses (118,119). Most studies which link IBS to LPS focus on levels of LPS in blood, and much of the information regarding levels of LPS in stool is conflicting. Although LPS can cause diarrhea through endotoxemia (120), the only consistent reports we could find linking IBS to fecal LPS levels indicated those with constipation were more likely to have elevated fecal LPS (64,121–123). We anticipate LPS-induced inflammation is generally triggered via toll-like receptors (TLRs), primarily via TLR4 of the intestinal epithelium, but also via TLRs located throughout the body and organs such as the liver (124).

The ORs for IBS-C highlight one more notable finding: for two scores, individuals with IBS-C are less likely to present with “Not Optimal” scores for *ButyrateProductionPathways* and *SulfideProductionPathways*). These values are inverted from IBS-D, and this is best explained in the context of IBS-C individuals being more likely to present with “Not Optimal” *MethanogenesisPathways* scores, which corresponds to increased methane produced from SCFAs (Table 9). Methane production in the gut is well-documented and correlated with IBS-C (66,125). Recently, it has been shown to affect gut transit time and intestinal peristaltic activity, eventually leading to constipation (125). It is also naturally correlated with bloating and flatulence (126). In light of increased methane among those with IBS-C, it is not surprising for IBS-C individuals to have increased butyrate production (i.e., reduced likelihood of “Not Optimal” scores for *ButyrateProductionPathways*), because butyrate is a SCFA that directly or indirectly (i.e., via acetate) promotes both methanogenesis (127) and constipation (128)). We can also expect a trend for IBS-C individuals to generally present with decreased levels of sulfide (i.e., decreased likelihood of “Not Optimal” scores for *SulfideProductionPathways*), because sulfate reduction often restricts methanogenesis via several pathways (129), with some exceptions (130,131). Furthermore, IBS studies often report sulfide as a driver of IBS-D and methane as a driver of IBS-C (66,132).

### Application of gut scores to dietary modifications

The implementation of dietary modifications based on these scores is beyond the scope of this manuscript, but we wanted to include some examples of such recommendations to benefit those interested in the application of biomarkers to wellness. First and foremost, we should be reminded that both dietary and medical clinicians utilize a wide variety of diagnostic tests, diagnostic aids, and “early health insights”, and after considering all of the available information, clinicians implement their own dietary or personalized treatment protocols in order to address any area deemed to be “Not Optimal” for an individual. Within our (direct-to-consumer) wellness platform, scores of “Not Optimal” receive dietary recommendations for individuals to consider. For example, *UricAcidProductionPathways* is implemented such that individuals with a “Not Optimal” score are encouraged to avoid foods high in purines, such as scallops, clams, and lobster, effectively reducing substrates for the microbial production of uric acid (133,134). Additionally, an increase in the consumption of polyphenols known to modulate both host and microbial uric acid metabolism are encouraged through both diet and supplementation. These can include citrus flavonoids like naringin, chlorogenic acid, fisetin, quercetin, and ferulic acid (135–140). For “Not Optimal” *GABAProductionPathways*, scores our wellness program encourages substrates and cofactors, such as _L_-glutamate and vitamin B6, known to be involved in GABA production (141–145). Probiotics observed to produce GABA are also supplemented (146–148). Excessive sulfide and methane gas, as indicated by “Not Optimal” *SulfideProductionPathways* and *MethanogenesisPathways* scores, are addressed through the restriction of sulfur containing substrates known to increase the microbial production of sulfide gas; the restriction of substrates like carnitine, choline, and fructooligosaccharides known to contribute to the production of microbial methane gas; as well as the introduction of polyphenols, prebiotics, and probiotics shown to reduce both the abundance and activity of gas producing microbes (149–159).

The impact these and other personalized interventions have on gastrointestinal function and mental health are currently being investigated (ClinicalTrials.gov ID’s NCT06190184; NCT05465629). The clinical application and assessment of Viome’s Precision Nutrition Plan were presented in an earlier manuscript that focused on pilot interventional studies (160). Each of these studies demonstrated significant improvements across various conditions as determined using clinically-validated surveys: IBS-SSS for IBS (161), PHQ9 for depression (162), GAD7 for anxiety (163), and an internally-developed risk score for type 2 diabetes (164).

### Clinical relevance of transcriptome-based gut pathway scores

Viome’s underlying metatranscriptome laboratory assay is an unbiased RNA detection method for the entire range of microbial activities (i.e. RNA molecules from any organism) in the gut. The assay itself provides an extremely broad set of RNA molecular biomarkers that can used for multiple downstream clinical applications, such as (a) diagnostic tests designed for identifying current disease state, leading to curative interventional procedures, (b) early detection tests (diagnostic aids), designed to detect subclinical disease leading to definitive diagnostic testing, or (c) “early health insights” designed to determine the activity of known harmful pathways, leading to preventive nutritional (or other lifestyle) modifications.

The metatranscriptome-based gut pathway scores presented in this paper are designed for “early health insights” of harmful molecular activities that could potentially lead to future disease state. For example, excess methane or sulfide pathway activities can lead to inflammation, which in turn can lead to multiple gut diseases. The intent behind the pathway scores presented here is to encourage changes to nutritional intake that lead to bringing these pathways more in line with the normal population distributions.

In other research, Viome has also developed early detection tests (diagnostic aids) (165) designed to detect disease-associated molecular pathways with high specificity and sensitivity, for the purpose of recommending further definitive diagnostic testing. Furthermore, Viome is developing early detection tests to distinguish between one DGBI and another, e.g. between IBS and IBD. It is indeed possible to use the scores presented in this paper as inputs into such early detection tests, but that is a topic for future research. For clarity, all early detection and diagnostic testing is outside the scope of this paper.

With regards to interpretation of ORs, while clinical standards vary, values above 2.0 are generally considered to represent a strong association that carries clinical relevance with actionability in standard clinical practice (166–168). In this context, ORs are common statistics that may help to guide decisions related to medical treatment care.

### Gut Microbiota in Other Diseases

Although the current study focuses on validating stool-based functional scores in the context of IBS and its subtypes, microbial functions contribute to a range of conditions beyond the gastrointestinal tract (i.e., systemic inflammation, autoimmune diseases, cardiometabolic disorders, and various neurological and psychiatric conditions). One area of interest includes the connections between gut microbiota and non-alcoholic fatty liver disease (NAFLD). Studies indicate that individuals with NAFLD exhibit distinct gut microbiomes with altered functions involving bile acid metabolism (169), choline degradation (170), and endotoxin production (171). Elevated gut-derived endotoxins, such as lipopolysaccharides (LPS), can enter the liver via the portal vein (which carries blood directly from the intestinal tract), triggering hepatic inflammation—a key factor in the progression from NAFLD to non-alcoholic steatohepatitis (NASH) (170,172). In this context, inflammation-related microbial functions may provide early molecular indicators of liver dysfunction, preceding clinical symptoms. Importantly, researchers have also demonstrated an ability to improve these condition-related indicators, underscoring that dysbiotic microbiomes can be therapeutically addressed (173–176). Advancing the ability to assess human microbiomes may afford improved ability to target specific microbial pathways implicated in disease pathogenesis. Future research should continue to expand across diverse clinical populations and disease contexts, integrating anthropomorphic and clinical metadata with multi-omics approaches intended to disentangle complex host-microbe interactions and identify actionable microbial signatures.

### Limitations

Several limitations exist for this study. The analysis includes case/control comparisons as part of its validation but may not address all potential confounding factors. The current analyses cannot establish causality due to the composition of scores, which likely consist of KO features comprising both causal and consequential elements. Future studies will be conducted to delineate causal features through prospective interventional trials. Metabolite levels could not be assessed in the current study, but we are planning to accomplish these measurements in future studies. While our cohorts are extensive and encompass various demographics, they may not entirely mirror every demographic group in the USA or other nations. Additionally, the metadata labels rely on self-reported data, which may be prone to misreporting; however, given the very large size of our cohorts, any such discrepancies are likely to be insignificant. Furthermore, incorporation of the independent and clinically validated Rome IV Diagnostic Questionnaire should mitigate concerns related to ORs for self-reported IBS phenotypes.

## Conclusions

To conclude, we summarize here the novel methods, findings, and some implications for current applications and future research.

Using stool metatranscriptomic data from a large adult population (*n* = 9,350), we have developed a suite of gut microbiome functional pathway scores that represent the biochemical activity of well-known processes in the gastrointestinal tract. The process for developing these pathway scores includes 5 iterative steps: domain exploration, metadata curation, signal definition, feature selection and pathway activity quantification. We employ a normative approach to determining the optimal and non-optimal segments of the score distributions within the development cohort. To verify that these pathways are consistent with their intended purpose, we confirm that the pathways are differentially active in a case/control analysis of IBS and subtypes within a large independent adult population (*n* = 71,220). Furthermore, we compute the odds that a non-optimal pathway activity of each of the scores is associated with IBS or its subtypes.

The main finding in this paper is that there is a significantly higher odds of IBS and subtypes for several of the scores, regardless of whether the IBS phenotype was determined via Rome IV or via self-reporting. In particular, individuals with not optimal *GABAProductionPathways* have higher odds of IBS and its subtypes (IBS-C, IBS-D, and IBS-M). In addition, individuals with not optimal *LacticAcidProduction-Pathways, OxalateMetabolismPathways*, or *Sulfide-ProductionPathways* have higher odds of IBS-D. Finally, individuals with not optimal *MethanogenesisPathways* have higher odds of IBS-C, while at the same time, individuals with not optimal *ButyrateProductionPathways* or *SulfideProductionPathways* have a lower odds of IBS-C.

Our findings demonstrate the use of metatranscriptomics for wellness applications like early health insights and nutritional recommendations, which we have previously implemented within a large-scale health application (160).

Supplementary Figure 1: Landscape of score distribution; Supplementary Table 1: Control definitions for each score; Supplementary Table 2: ORs and Confidence Intervals shown in Figure 3; Supplementary Table 3: ORsand Confidence Intervals shown in Figure 4; Supplementary Figure 2: Venn diagram for genera of significantly increased species across self-reported IBS phenotypes; Supplementary Figure 3: Venn diagram for genera of significantly decreased species across self-reported IBS phenotypes; Supplementary Figure 4: Venn diagram for genera of significantly increased species across Rome IV-defined IBS phenotypes; Supplementary Figure 5: Venn diagram for genera of significantly decreased species across Rome IV-defined IBS phenotypes.

## Declarations

## Acknowledgements

We would like to thank Angel Janevski for leading data and analytics, which is responsible for computing scores and recommendations on a regular basis.

## Author Contributions

Conceptualization, E.P., G.B.; Data curation, E.P., A.G., M.M.; Formal analysis, E.P., A.G., M.M., O.O.; Funding acquisition, M.V., G.B.;

Investigation, E.P., A.G., M.M., C.J., L.H., M.V., G.B.; Methodology, E.P., A.G., M.M.; Project administration, G.B.; Resources, G.B.; Software, E.P., A.G., M.M., O.O., L.H.; Supervision, G.B.; Validation, C.J.; Visualization, E.P., A.G., M.M.; Writing – original draft, E.P., A.G., M.M., G.B.; Writing – review & editing, E.P., A.G., M.M., C.J., L.H., G.A., M.V., R.W., G.B.

## Availability of Data and Materials

The data that support the findings of this study are from Viome Life Sciences Inc but restrictions apply to the availability of these data, which were used under the terms of an agreement for the current study, and so are not publicly available. Data in the form of summary statistics are however available for non-commercial purposes through a Data Transfer Agreement from the author Guru Banavar upon reasonable request via: https://www.viomelifesciences.com/data-access.

## Competing Interests

All authors are stockholders and either employees or paid advisors of Viome Inc, a commercial for-profit company.

## Consent for publication

Not applicable.

## Ethics approval and consent to participate

All customers were at least 18 years old at the time of sample collection. All customers were informed, and they consented to their data being used for research purposes, as part of the sign-up process for Viome services. At Viome Life Sciences, we assess each research project to determine whether it needs to be reviewed by an Institutional Review Board (IRB). Based on the US Health and Human Services CFR 46.104, section 4(II), the Ethics Program Director determined that this research is exempt from IRB review. This research project solely uses retrospective data from Viome Life Sciences’ customers. All study data are de-identified; data analysis team members have no access to personally identifiable information.

## Funding

No funding was received outside of Viome.

## Extended Data

**Extended Data Figure 1.**
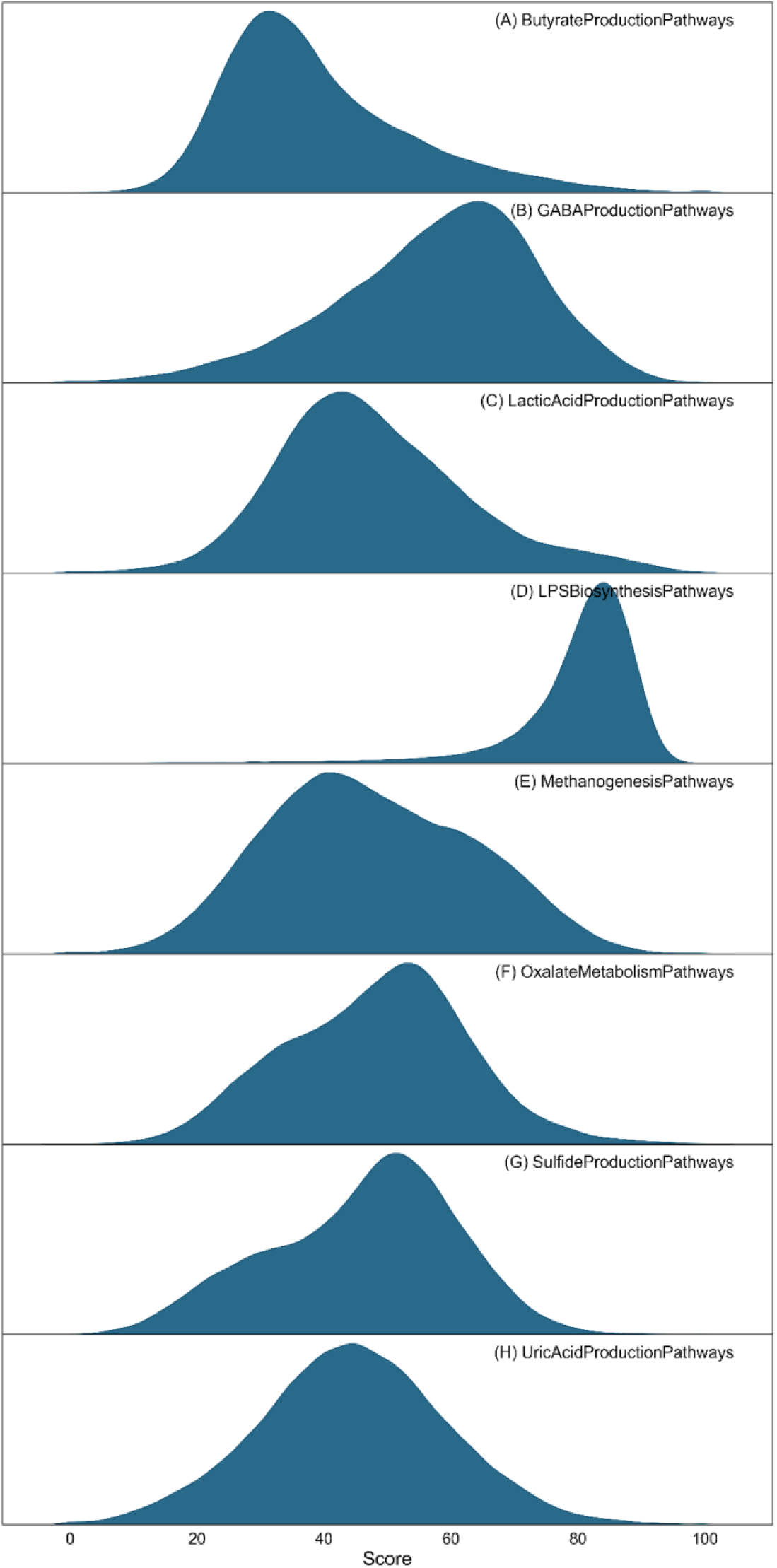
Landscape of score distribution. The distribution of eight functional pathway scores (A-H) in an independent cohort (*n* = 71,220 samples). The distribution of each pathway score across the entire cohort is expected to follow a Gaussian normal distribution, where a high score is “Good”. Some scores deviate significantly from this normal distribution, which can be attributed to missing molecular signatures in the reference cohort, resulting in extremely high or low scores. Scores exceeding 100 or falling below 0 were clipped.

**Extended Data Figure 2.**
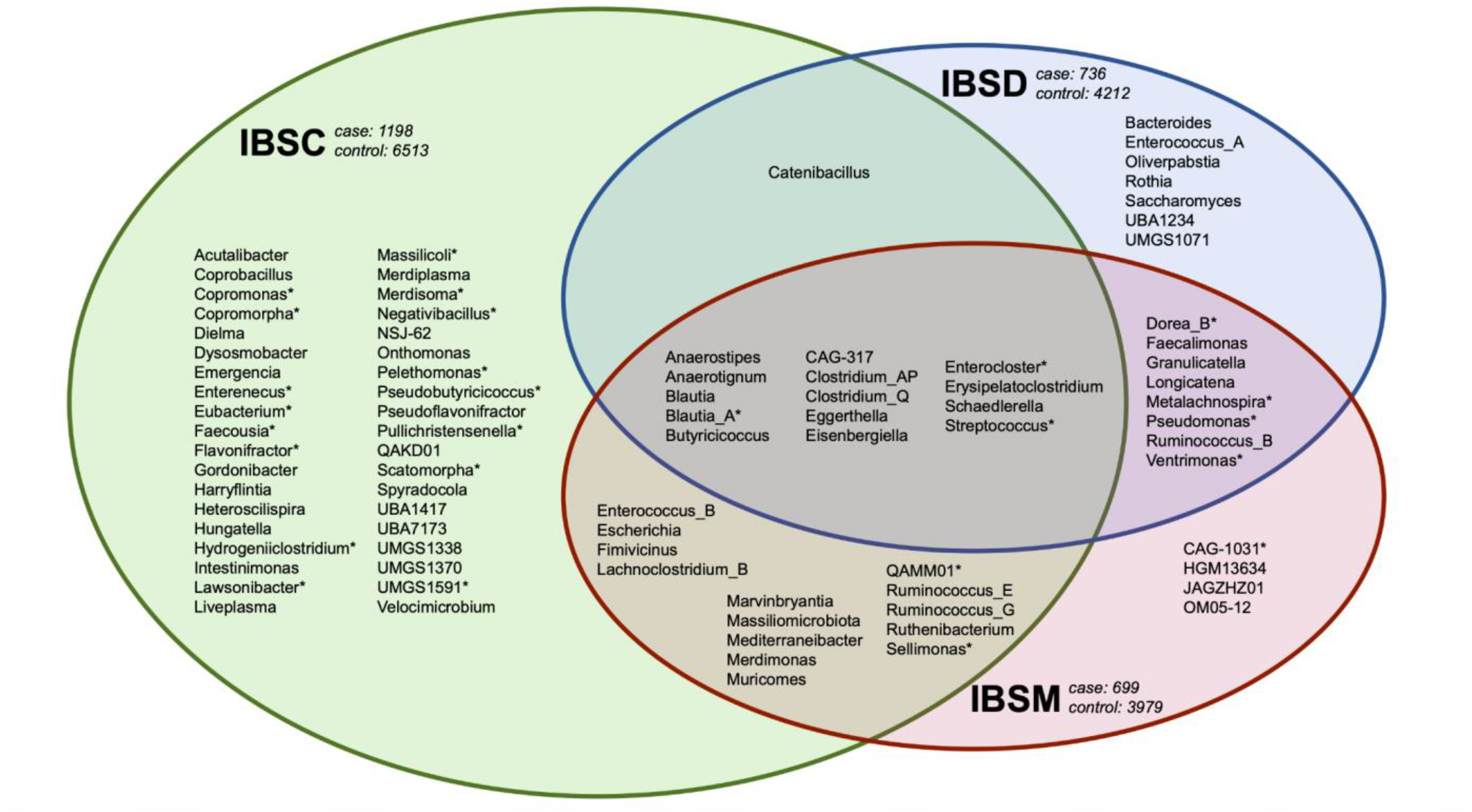
Genera of significantly increased species (differentially expressed) between self-reported IBS phenotypes and individuals with no comorbidities. Controls are defined as individuals with zero self-reported comorbidities. At the highest level, cases and controls were matched 1:5 for “IBS-C” (green; *n* = 1,198), “IBS-D” (blue; *n* = 736), and “IBS-M” (red; *n* = 699). All significantly increased species (p ≤ 0.05 and log “Fold Change” ≥ 0.6) were combined into respective genera before constructing the Venn Diagram. All genera with differential expression consistent to Rome IV-determined IBS phenotypes (Extended Data Figure 4) are indicated with an asterisk (*).

**Extended Data Figure 3.**
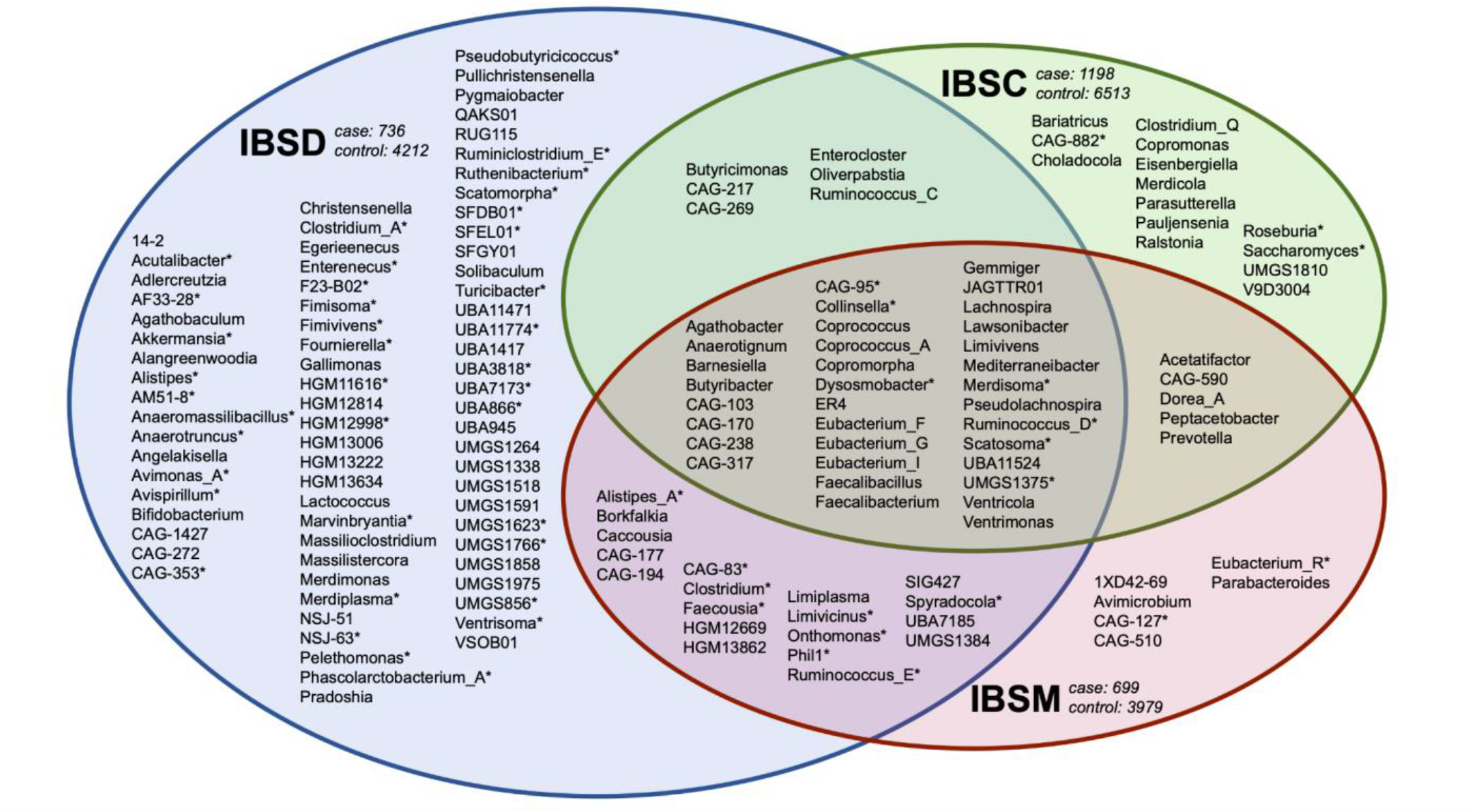
Genera of significantly decreased species (differentially expressed) between self-reported IBS phenotypes and individuals with no comorbidities. Controls are defined as individuals with zero self-reported comorbidities. At the highest level, cases and controls were matched 1:5 for “IBS-C” (green; *n* = 1,198), “IBS-D” (blue; *n* = 736), and “IBS-M” (red; *n* = 699). All significantly decreased species (p ≤ 0.05 and log “Fold Change” ≤ −0.6) were combined into respective genera before constructing the Venn Diagram. All genera with differential expression consistent to Rome IV-determined IBS phenotypes (Extended Data Figure 5) are indicated with an asterisk (*).

**Extended Data Figure 4.**
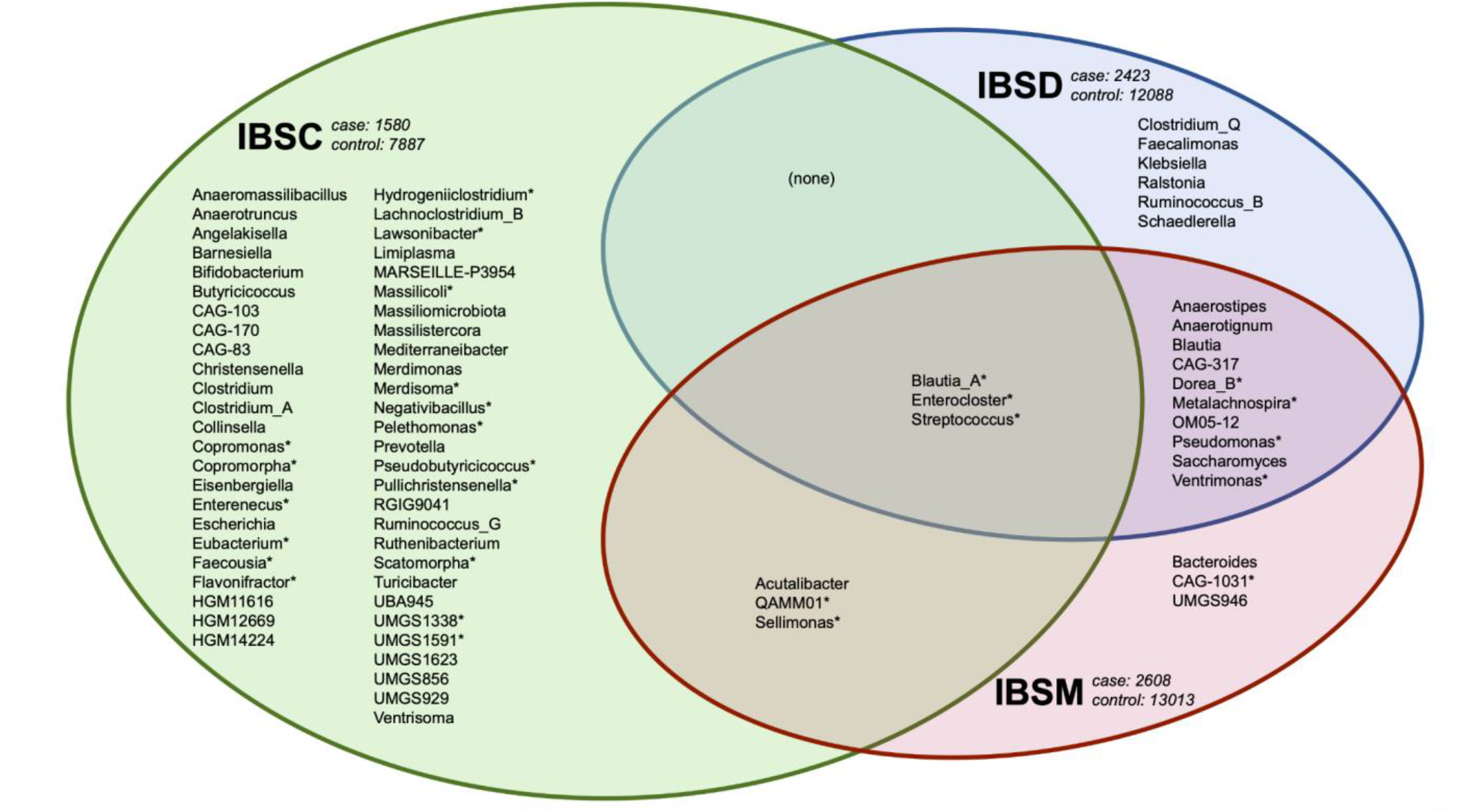
Genera of significantly increased species (differentially expressed) between Rome IV-defined IBS phenotypes and individuals with no comorbidities. Controls are defined as individuals with zero self-reported comorbidities. At the highest level, cases and controls were matched 1:5 for “IBS-C” (green; *n* = 1,198), “IBS-D” (blue; *n* = 736), and “IBS-M” (red; *n* = 699). All significantly increased species (p ≤ 0.05 and log “Fold Change” ≥ 0.6) were combined into respective genera before constructing the Venn Diagram. All genera with differential expression consistent to self-reported IBS phenotypes (Extended Data Figure 2) are indicated with an asterisk (*).

**Extended Data Figure 5.**
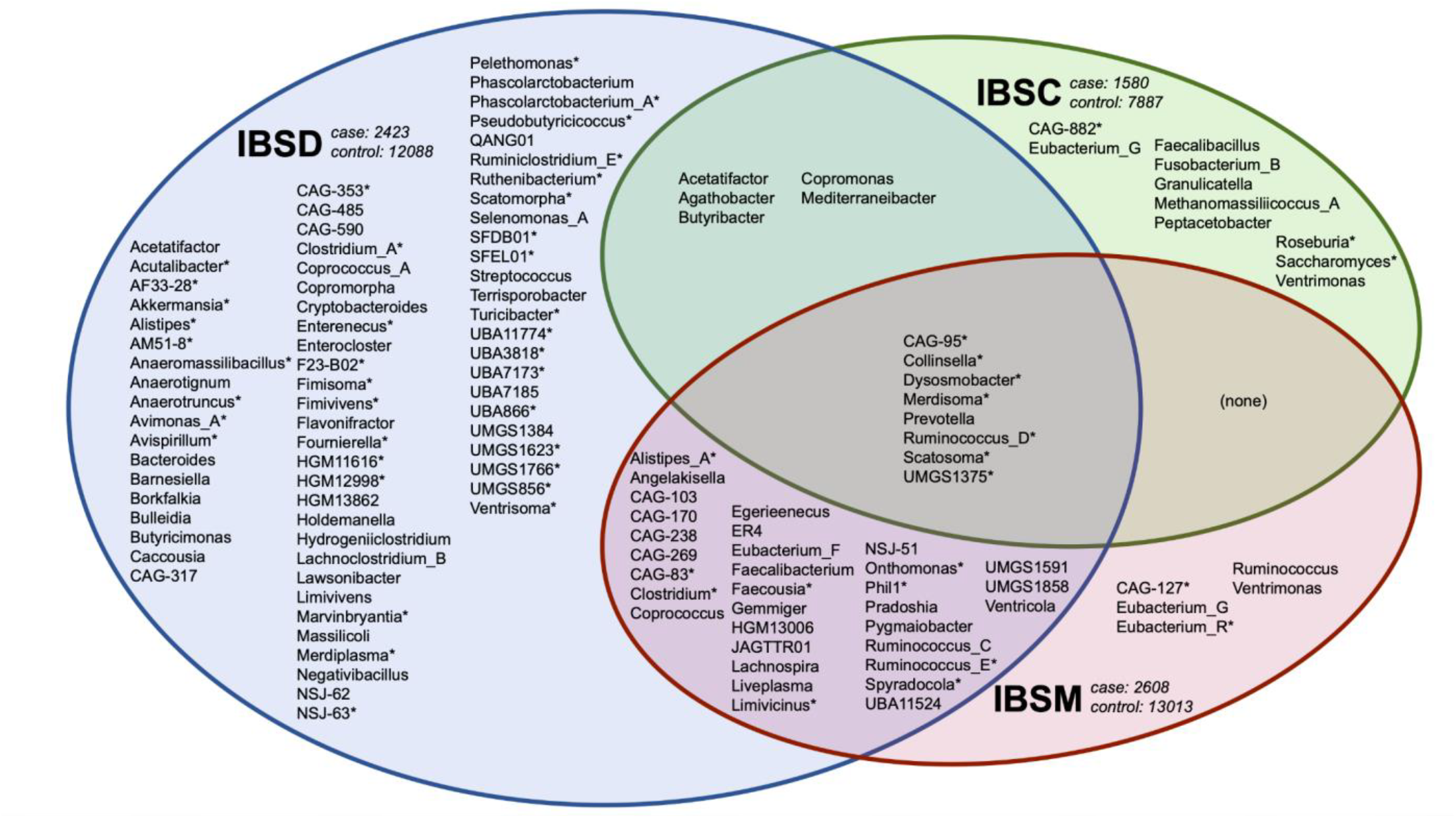
Genera of significantly decreased species (differentially expressed) between Rome IV-defined IBS phenotypes and individuals with no comorbidities. Controls are defined as individuals with zero self-reported comorbidities. At the highest level, cases and controls were matched 1:5 for “IBS-C” (green; *n* = 1,198), “IBS-D” (blue; *n* = 736), and “IBS-M” (red; *n* = 699). All significantly increased species (p ≤ 0.05 and log “Fold Change” ≥ 0.6) were combined into respective genera before constructing the Venn Diagram. All genera with differential expression consistent to self-reported IBS phenotypes (Extended Data Figure 3) are indicated with an asterisk (*).

**Extended Data Table 1.**
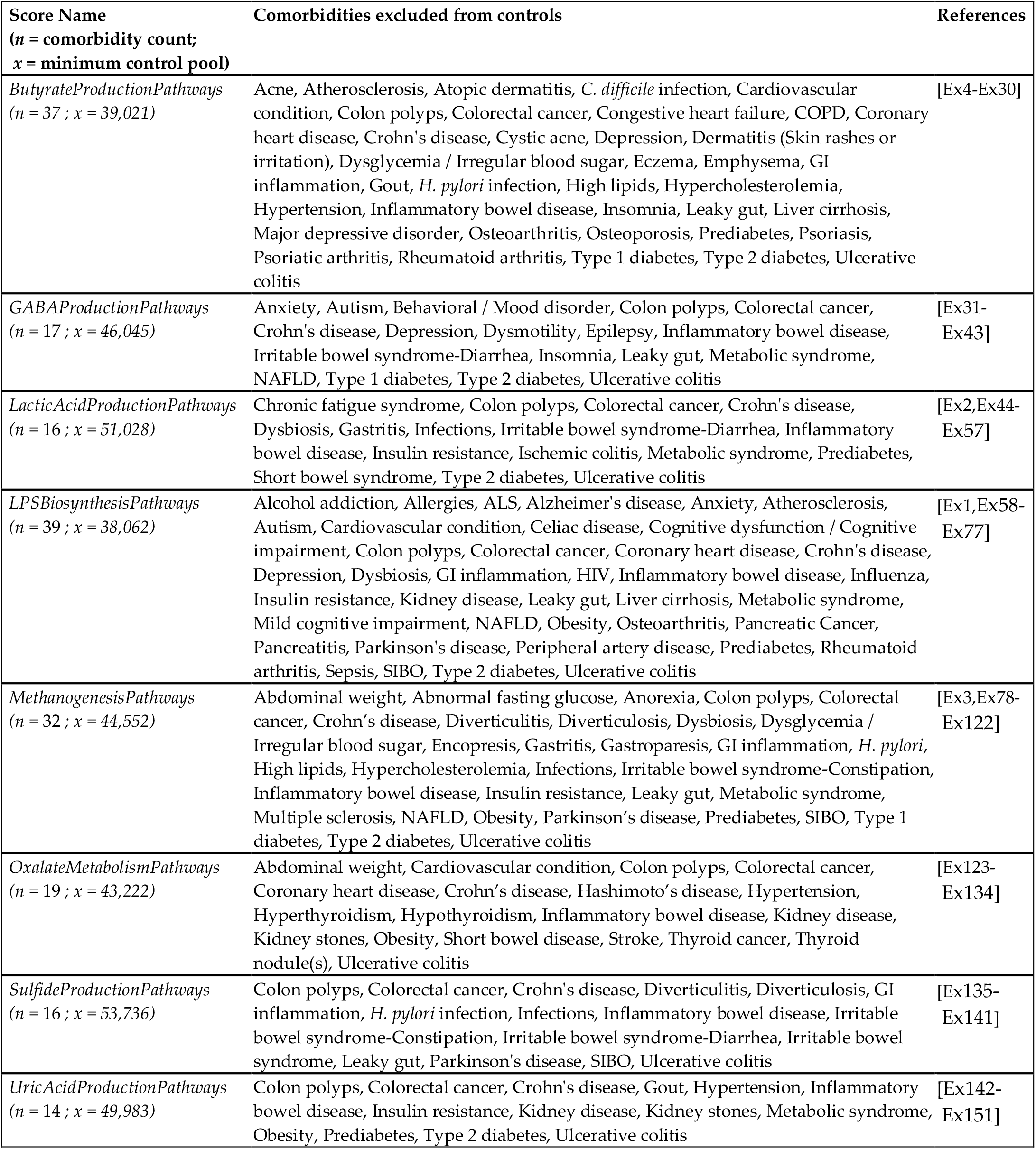
Control definitions for each score. The definition of “controls” shown in Figure 3 is different for each score, where all diseases listed for each score were removed from the control pool, along with anyone who reported IBS or its subtypes. The count of comorbidities (*n*) is shown alongside the minimum size of the control pool (*x*) available for 1:5 matching, which was determined by subtracting the number of excluded cases (plus all IBS cases) from the size of the validation cohort (*n* = 71,220). For odds ratios based on Rome IV criteria, any positive IBS classification was also excluded from controls.

| Score Name<br>(*n* = comorbidity count;<br>*x* = minimum control pool) | Comorbidities excluded from controls | References |
| --- | --- | --- |
| *ButyrateProductionPathways*<br>(*n* = 37 ; *x* = 39,021) | Acne, Atherosclerosis, Atopic dermatitis, *C. difficile* infection, Cardiovascular condition, Colon polyps, Colorectal cancer, Congestive heart failure, COPD, Coronary heart disease, Crohn's disease, Cystic acne, Depression, Dermatitis (Skin rashes or irritation), Dysglycemia / Irregular blood sugar, Eczema, Emphysema, GI inflammation, Gout, *H. pylori* infection, High lipids, Hypercholesterolemia, Hypertension, Inflammatory bowel disease, Insomnia, Leaky gut, Liver cirrhosis, Major depressive disorder, Osteoarthritis, Osteoporosis, Prediabetes, Psoriasis, Psoriatic arthritis, Rheumatoid arthritis, Type 1 diabetes, Type 2 diabetes, Ulcerative colitis | [Ex4-Ex30] |
| *GABAProductionPathways*<br>(*n* = 17 ; *x* = 46,045) | Anxiety, Autism, Behavioral / Mood disorder, Colon polyps, Colorectal cancer, Crohn's disease, Depression, Dysmotility, Epilepsy, Inflammatory bowel disease, Irritable bowel syndrome-Diarrhea, Insomnia, Leaky gut, Metabolic syndrome, NAFLD, Type 1 diabetes, Type 2 diabetes, Ulcerative colitis | [Ex31-Ex43] |
| *LacticAcidProductionPathways*<br>(*n* = 16 ; *x* = 51,028) | Chronic fatigue syndrome, Colon polyps, Colorectal cancer, Crohn's disease, Dysbiosis, Gastritis, Infections, Irritable bowel syndrome-Diarrhea, Inflammatory bowel disease, Insulin resistance, Ischemic colitis, Metabolic syndrome, Prediabetes, Short bowel syndrome, Type 2 diabetes, Ulcerative colitis | [Ex2,Ex44-Ex57] |
| *LPSBiosynthesisPathways*<br>(*n* = 39 ; *x* = 38,062) | Alcohol addiction, Allergies, ALS, Alzheimer's disease, Anxiety, Atherosclerosis, Autism, Cardiovascular condition, Celiac disease, Cognitive dysfunction / Cognitive impairment, Colon polyps, Colorectal cancer, Coronary heart disease, Crohn's disease, Depression, Dysbiosis, GI inflammation, HIV, Inflammatory bowel disease, Influenza, Insulin resistance, Kidney disease, Leaky gut, Liver cirrhosis, Metabolic syndrome, Mild cognitive impairment, NAFLD, Obesity, Osteoarthritis, Pancreatic Cancer, Pancreatitis, Parkinson's disease, Peripheral artery disease, Prediabetes, Rheumatoid arthritis, Sepsis, SIBO, Type 2 diabetes, Ulcerative colitis | [Ex1,Ex58-Ex77] |
| *MethanogenesisPathways*<br>(*n* = 32 ; *x* = 44,552) | Abdominal weight, Abnormal fasting glucose, Anorexia, Colon polyps, Colorectal cancer, Crohn's disease, Diverticulitis, Diverticulosis, Dysbiosis, Dysglycemia / Irregular blood sugar, Encopresis, Gastritis, Gastroparesis, GI inflammation, *H. pylori*, High lipids, Hypercholesterolemia, Infections, Irritable bowel syndrome-Constipation, Inflammatory bowel disease, Insulin resistance, Leaky gut, Metabolic syndrome, Multiple sclerosis, NAFLD, Obesity, Parkinson's disease, Prediabetes, SIBO, Type 1 diabetes, Type 2 diabetes, Ulcerative colitis | [Ex3,Ex78-Ex122] |
| *OxalateMetabolismPathways*<br>(*n* = 19 ; *x* = 43,222) | Abdominal weight, Cardiovascular condition, Colon polyps, Colorectal cancer, Coronary heart disease, Crohn's disease, Hashimoto's disease, Hypertension, Hyperthyroidism, Hypothyroidism, Inflammatory bowel disease, Kidney disease, Kidney stones, Obesity, Short bowel disease, Stroke, Thyroid cancer, Thyroid nodule(s), Ulcerative colitis | [Ex123-Ex134] |
| *SulfideProductionPathways*<br>(*n* = 16 ; *x* = 53,736) | Colon polyps, Colorectal cancer, Crohn's disease, Diverticulitis, Diverticulosis, GI inflammation, *H. pylori* infection, Infections, Inflammatory bowel disease, Irritable bowel syndrome-Constipation, Irritable bowel syndrome-Diarrhea, Irritable bowel syndrome, Leaky gut, Parkinson's disease, SIBO, Ulcerative colitis | [Ex135-Ex141] |
| *UricAcidProductionPathways*<br>(*n* = 14 ; *x* = 49,983) | Colon polyps, Colorectal cancer, Crohn's disease, Gout, Hypertension, Inflammatory bowel disease, Insulin resistance, Kidney disease, Kidney stones, Metabolic syndrome, Obesity, Prediabetes, Type 2 diabetes, Ulcerative colitis | [Ex142-Ex151] |

**Extended Data Table 2.** Odds Ratio and Confidence Intervals shown in Figure 3. IBS cases are the composite of IBS and all IBS subtypes while “control” definitions are score-specific (Extended Data Table 1). Values represent Odds Ratios. Significance levels of the p-values are shown in superscript, with 95% Confidence Interval in parentheses. P ≤ 0.05 (^*)^; p ≤ 0.01 (**); p ≤ 0.001 (***); not significant (ns).

| Score Name | IBS<br>(max $n = 6,373$) | IBS-C<br>(max $n = 1,732$) | IBS-D<br>(max $n = 1,006$) | IBS-M<br>(max $n = 1,091$) |
| --- | --- | --- | --- | --- |
| *ButyrateProductionPathways* | 1.3*** (1.2 - 1.5) | 0.8* (0.7 - 0.9) | 2.6*** (2.1 - 3.3) | 1.5*** (1.2 - 1.9) |
| *GABAProductionPathways* | 1.7*** (1.5 - 1.8) | 2.1*** (1.8 - 2.6) | 1.6*** (1.3 - 2.0) | 1.5*** (1.2 - 1.9) |
| *LacticAcidProductionPathways* | 1.4*** (1.3 - 1.5) | 0.9<sup>ns</sup> (0.8 - 1.1) | 2.3*** (1.8 - 2.9) | 1.2<sup>ns</sup> (1.0 - 1.5) |
| *LPSBiosynthesisPathways* | 1.2** (1.1 - 1.3) | 1.3** (1.1 - 1.5) | 0.9<sup>ns</sup> (0.7 - 1.1) | 1.1<sup>ns</sup> (0.9 - 1.4) |
| *MethanogenesisPathways* | 1.2*** (1.1 - 1.4) | 1.6*** (1.4 - 2.0) | 1.2<sup>ns</sup> (1.0 - 1.5) | 1.3* (1.0 - 1.5) |
| *OxalateMetabolismPathways* | 1.3*** (1.2 - 1.4) | 0.9<sup>ns</sup> (0.7 - 1.0) | 2.1*** (1.6 - 2.6) | 1.2* (1.0 - 1.5) |
| *SulfideProductionPathways* | 1.1<sup>ns</sup> (1.0 - 1.2) | 0.7*** (0.6 - 0.9) | 1.8*** (1.5 - 2.3) | 1.1<sup>ns</sup> (0.9 - 1.3) |
| *UricAcidProductionPathways* | 1.5*** (1.4 - 1.6) | 1.0<sup>ns</sup> (0.8 - 1.1) | 2.3*** (1.8 - 2.9) | 1.3* (1.1 - 1.6) |

**Extended Data Table 3.**
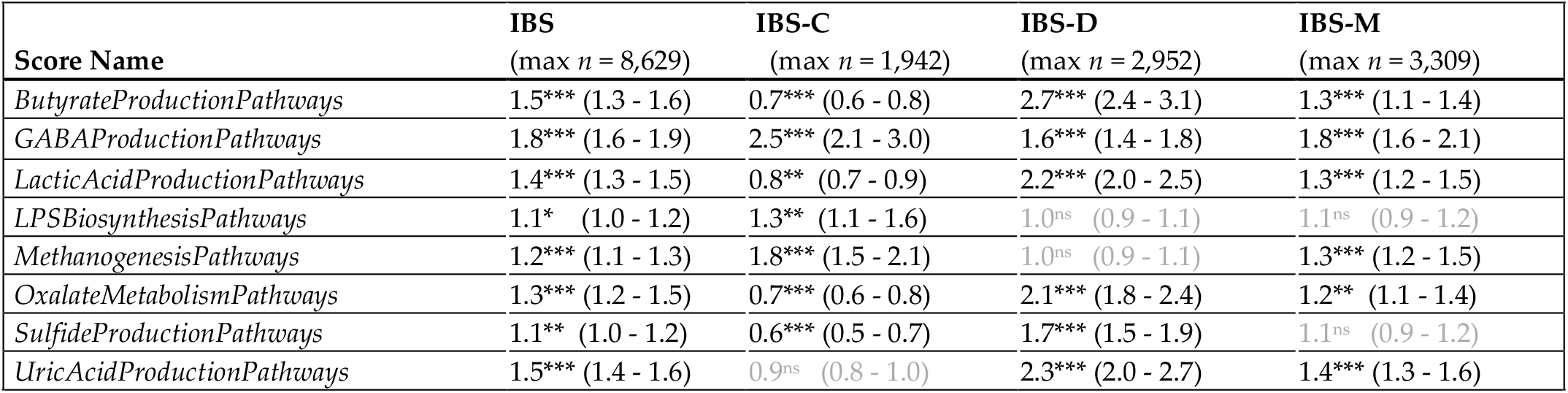
Odds Ratio and Confidence Intervals shown in Figure 4. IBS cases are the composite of any IBS subtype classification, and cases for each IBS subtype classification are defined by the Rome IV Diagnostic Questionnaire. “Control” definitions are score-specific (Extended Data Table 1). Values represent Odds Ratios. Significance levels of the p-values are shown in superscript, with 95% Confidence Interval in parentheses. p ≤ 0.05 (*); p ≤ 0.01 (**); p ≤ 0.001 (***); not significant (ns).

## Notes

### Summary of Updates

Removing duplicate references, revising table citations, and moving "supplementary references" to a section with the supplementary materials.

